# Cyclin binding Cy motifs have multiple activities in the initiation of DNA replication

**DOI:** 10.1101/681668

**Authors:** Manzar Hossain, Kuhulika Bhalla, Bruce Stillman

## Abstract

The initiation of DNA replication involves the cell cycle-dependent assembly and disassembly of protein complexes, including the Origin Recognition Complex (ORC) and CDC6 AAA+ ATPases. We report that multiple short, linear protein motifs (SLiMs) within intrinsically disordered regions in ORC1 and CDC6, including Cyclin-binding (Cy) motifs, mediate Cyclin-CDK dependent and independent protein-protein interactions, conditional on cell cycle phase. The ORC1 Cy motif mediates an auto-regulatory self-interaction, and the same Cy motif prevents CDC6 binding to ORC1 in mitosis, but then facilitates the destruction of ORC1 in S phase. In contrast, in G1, the CDC6 Cy motif promotes ORC1-CDC6 interaction independent of Cyclin-CDK protein phosphorylation. CDC6 interaction with ORC also requires a basic region of ORC1 that in yeast mediates ORC-DNA interactions. We also demonstrate that protein phosphatase 1 binds directly to a SLiM in ORC1, causing de-phosphorylation upon mitotic exit. Thus, Cy-motifs have wider roles, functioning as a ligand and as a degron.

## Introduction

Comparative studies of gene regulation and control of molecular circuitries across different species has revealed that a larger genome size need not correlate with biological or proteomic complexity of the organism. What has emerged as the fundamental driver of this complexity is an increase in regulatory networks rather than absolute protein numbers (Hahn and Wray, 2002; Schad et al., 2011). Comparison of cell division cycle regulation in many model species exemplifies this observation – as even though core components of the machinery itself are conserved across eukaryotic evolutionary timeline, metazoan systems experience significantly more elaborate functional regulation, compared to their simpler eukaryotic counterparts such as budding yeast (O’Donnell et al., 2013). Advances in our understanding of higher eukaryotic interactomes indicate the presence of a myriad of short linear protein motifs (SLiMs) that mediate a cascade of interactions and/or functions. While all proteins contain various large structured domains to which function can be ascribed, many short regulatory motifs have been identified in proteins, frequently within intrinsically disordered regions (IDR), perhaps to allow for greater combinatorial or conditional regulation by harnessing the region’s structural plasticity (Russell and Gibson, 2008). SLiMs in proteins have been broadly categorized based on discrete functional output such as mediating protein-protein interactions (e.g. SH3 ligands), subcellular localization (e.g. nuclear export signal), enzyme modification (e.g. Cyclin Dependent Kinase [CDK] docking motif), protein targeting for degradation (e.g. APC/C degron) and post-translational modifications (e.g. phosphorylation consensus sites) (Van Roey and Davey, 2015; Van Roey et al., 2014). Even though ∼85% of SLiMs occur in IDRs and several DNA replication initiator proteins harbor regions predicted to be intrinsically disordered, their functions in DNA replication are not understood.

In the budding yeast, DNA replication initiates at multiple origins throughout S phase, following assembly at every potential origin of a pre-Replicative Complex (pre-RC) that requires the six subunit Origin Recognition Complex (ORC), Cdc6, Cdt1 and two hetero-hexameric (Mcm2-7) complexes during the G1 phase of the cell cycle (Bell and Labib, 2016; Parker et al., 2017). In contrast, many questions remain about pre-RC assembly in human cells. For example, it is not known when CDC6 binds to ORC owing to the fact that ORC1, the largest subunit of ORC, is ubiquitinated and targeted for degradation at the onset of S phase and then resynthesized in late G2 and binds chromosomes as cells enter into mitosis (DePamphilis, 2003; Kara et al., 2015; Kreitz et al., 2001; Mendez et al., 2002; Ohta et al., 2003; Tatsumi et al., 2003). Human CDC6 on the other hand undergoes proteolytic degradation mediated by the APC^CDH1^ upon mitotic exit and in early G1 phase, to then be re-synthesized and stabilized by Cyclin E-CDK2 dependent phosphorylation, a cycle completely opposite that of *S. cerevisiae* Cdc6 (Cook et al., 2002; Coverley et al., 2002; Duursma and Agami, 2005; Mailand and Diffley, 2005). Thereafter, during S phase CDC6 is phosphorylated by Cyclin A-CDK2 and relocalized to the cytoplasm (Delmolino et al., 2001; Jiang et al., 1999; Petersen et al., 2000). This temporally restricted control and expression pattern of these key origin recognition and pre-RC components contributes to genome integrity by ensuring origins are licensed only once in G1 phase of the cell cycle. In *S. cerevisiae,* ORC recognizes sequence specific origins and all of its subunits, including Orc1, are stable and consequently ORC remains bound to origins of DNA replication for the entire cell cycle (Bell and Stillman, 1992; Diffley et al., 1994; Liang and Stillman, 1997; Tanaka et al., 1997). In budding yeast, however, Cdc6 levels vary as a consequence of SCF^Cdc4^ mediated degradation in S-phase, dependent on Cln-Cdc28 phosphorylation of Cdc6 (Figure 1A, bottom panel) (Perkins et al., 2001; Piatti et al., 1996) and the levels of yeast Cdc6 are reminiscent of human ORC1 (Figure 1A). Yeast Cdc6 is then re-synthesized during late G2 phase to enable binding to ORC following mitotic exit to form functional pre-RCs (Baum et al., 1998; Cocker et al., 1996; Drury et al., 2000; Piatti et al., 1995). Human ORC binds DNA in an apparent sequence-independent manner and with CDC6 and CDT1 can assembly MCM2-7 onto DNA *in vitro* (Vashee et al., 2003). In addition to their canonical roles in DNA replication, human ORC1 and CDC6 are involved directly in regulation of *CCNE1* gene expression in mid G1 phase to influence the decision of whether cells will proliferate or not (Hossain and Stillman, 2016).

**Figure 1.**
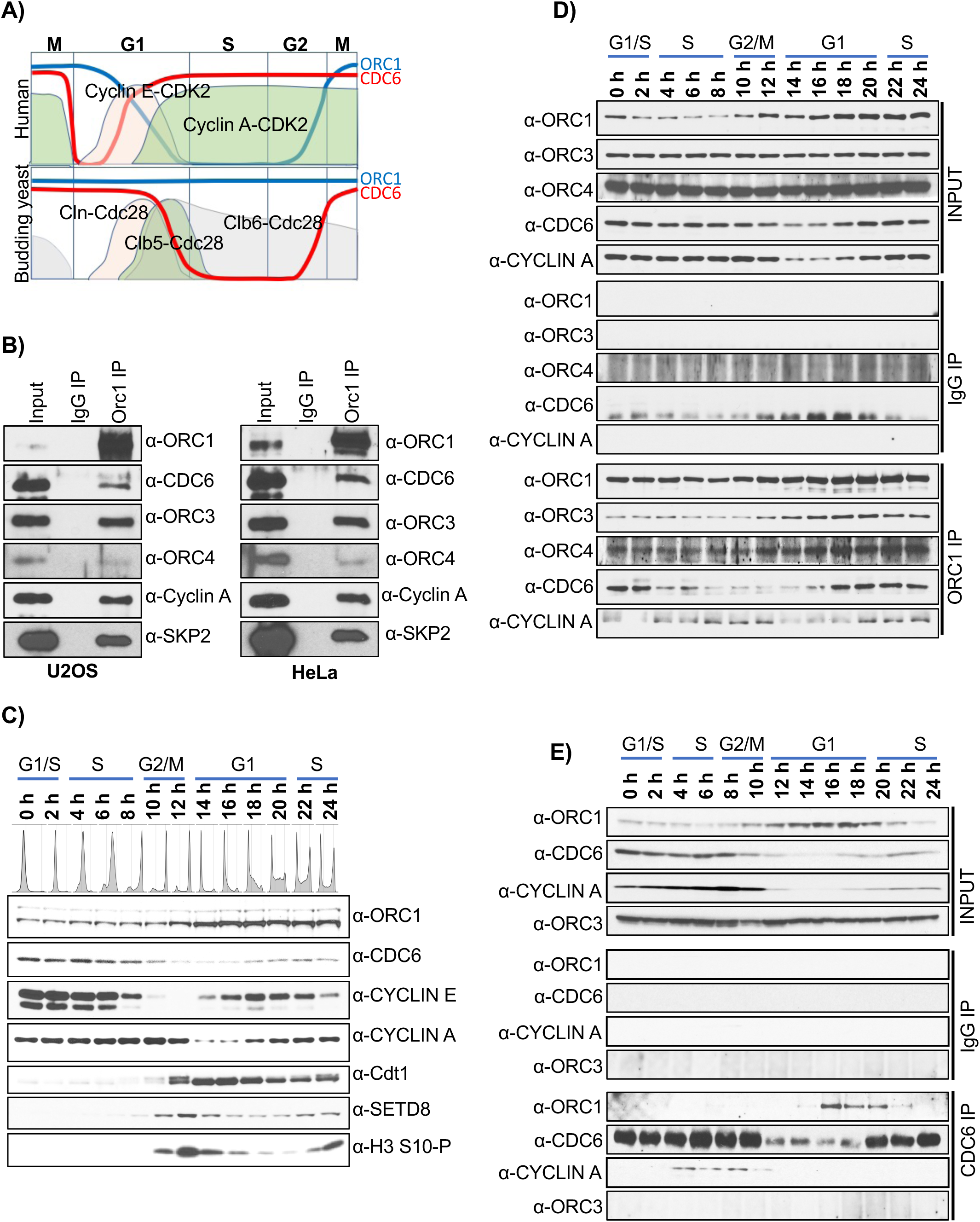
ORC1 and CDC6 proteins interact transiently at mid-G1 phase of the cell cycle. (A) Schematic of dynamic expression pattern of human and yeast ORC1 and CDC6 proteins across the cell division cycle with fluctuating levels of Cyclin-CDK kinases. (B) Immunoprecipitation of ORC1 from asynchronous U2OS (left panel) and HeLa (right panel) cell lysates showing interactions with CDC6, ORC3, ORC4, Cyclin A and SKP2 proteins. (C) Dynamic expression profile of pre-RC proteins such as ORC1, CDC6 and CDT1 along with cell cycle marker proteins (Cyclin E, Cyclin A, SETD8 and histone H3S10 phosphorylation) were studied by immunoblotting of extracts from double thymidine block and released synchronized HeLa cells. Flow cytometry profile of DNA content of synchronized HeLa cells at the indicated time in hours after release and corresponding phase of the cell cycle is indicated. (D-E) Double thymidine synchronized HeLa cell lysate at different time points were immunoprecipitated either with an ORC1 antibody (D) or a CDC6 antibody (E) and immunoblotted as indicated. The input and IgG IP denote loading control and mock IP in the experiment, respectively.

It has long been known that Cyclin-CDKs regulate the timing of pre-RC assembly and function, but whether Cyclin-CDKs have any direct effect on the interaction between human ORC and CDC6 is not known (Coverley et al., 2002; Li et al., 2004). Typically, the interaction with Cyclin-CDKs requires their substrates to harbor a specific Cyclin-CDK recognition motif (Cy motif with consensus R/KxL), which was first identified in the Cyclin-Cdk2 inhibitor proteins (CDKIs) (Adams et al., 1996; Takeda et al., 2001; Wohlschlegel et al., 2001). Since then it has been identified in numerous substrates of Cyclins A, B, D and E kinases, including ORC1 and CDC6 (Hemerly et al., 2009; Schulman et al., 1998; Wood and Endicott, 2018). Interestingly, the N-terminal regions in both of these proteins that harbor the Cy motifs and multiple CDK phosphorylation sites, are predicted to be intrinsically disordered (Figure S1). Notably, the entire N-terminal region (1-470 aa) of ORC1, including its extensive IDR (175-480 aa; Figure S1), was removed in order to determine the structure of the human ORC (ORC1-5) (Tocilj et al., 2017). The N-terminal regions of yeast Orc1 and Cdc6 homologs are also predicted to contain intrinsically disordered domains, though there is no apparent Cy motif in yeast Orc1 (Figure S1). In *S. cerevisiae*, Cdc6 and Orc6 contains a Cy motif that mediates Clb5-Cdc28 phosphorylation of ORC (Ord et al., 2019; Wilmes et al., 2004).

In this report, we show that the N-terminal disordered region of human ORC1 protein contains multiple SLiMs that perform varied regulatory functions during the cell cycle. Cyclin A-CDK2 mediated phosphorylation of ORC1 in mitosis is a newly identified regulatory step that blocks ORC1 binding to CDC6, thereby preventing premature pre-RC assembly. Protein phosphatase PP1 binds a short motif in the ORC1 IDR and partially dephosphorylates ORC1 upon mitotic exit, making ORC1 and CDC6 interaction permissive only during mid G1 phase of cell cycle. The Cy motif in the IDR of ORC1 also mediates an intramolecular interaction that is disrupted by CDC6 protein, a step that requires the CDC6 Cy motif. After pre-RC assembly in mid G1 phase, Cyclin A-CDK2 mediated phosphorylation of ORC1 and its recruitment of SCF^SKP2^ ubiquitin ligase causes the Cy motif dependent destruction of ORC1 as cells enter S phase. We propose that differences in the ORC-CDC6 interaction cycle between budding yeast and human cells are mediated, in large part, by the acquisition of multiple SLiMs in the IDRs of ORC1 and CDC6 proteins that are co-regulated by Cyclin-CDK kinase and protein phosphatases. These observations uncover expanded roles for the Cy motif, not only as an accelerator of CDK enzyme activity, but as a protein interaction ligand or module and a mediator of protein destruction, dependent on the stage of the cell division cycle.

## Results

### Dynamic interaction between ORC1 and CDC6 proteins during the human cell cycle

It is known that yeast Cdc6 and human CDC6 directly bind to yeast Orc1 and human ORC1, respectively (Hossain and Stillman, 2016; Liang et al., 1995; Saha et al., 1998). However, the yeast and human proteins have very different cell cycle expression profiles (Figure 1A). Because of this dynamic expression pattern of human ORC1 and CDC6 across the cell cycle, we investigated the biochemical interactions between these proteins. Using specifically designed antibodies against ORC1 and CDC6, ORC1 immunoprecipitation from asynchronous U2OS and HeLa cells showed co-precipitation of other ORC subunits, CDC6, Cyclin A and SKP2 (Figure 1B). To better understand the association between ORC1 and CDC6 across the mammalian cell cycle, HeLa cells grown in suspension were synchronized using double thymidine block. Synchrony of the cells collected at every 2-hour interval for 24 hours following release from second thymidine block was analyzed by flow cytometry and cell extracts were used for immunoblotting with known cell cycle marker proteins (Figure 1C). Cells were tightly synchronized with G1-S phase transition, S-phase, G2-M phase and G1 phase of next cell cycle occurring at around 0-2, 4-8, 10-12 and 14-20 hours, respectively. Thereafter cells entered the next S-phase and lost synchrony between 22-24 hours. In the synchronized cells, protein levels of Cyclin E and Cyclin A varied as expected, undergoing proteasome-mediated degradation at G1-S phase and upon mitotic exit, respectively. The protein levels of CDT1 and SETD8 were absent in S phase but were re-synthesized after mitosis, while histone H3S10 phosphorylation served as a marker for mitotic cells occurring at 12-hour time point (Figure 1C).

The interaction between ORC1 (Figure 1D) and CDC6 (Figure 1E) with each other and other proteins was investigated by immunoprecipitation. The level of ORC1 protein remained low until early G2-phase due to proteasome-mediated degradation at the G1/S transition. Upon exit from mitosis, ORC1 bound to ORC3 and ORC4 proteins and since both ORC3 and ORC4 proteins were expressed constitutively across the cell cycle, variation in their co-immunoprecipitation with ORC1 reflected ORC1 protein levels. Human ORC1 also interacted with cyclin A protein in S phase and mitosis when both were present. Interestingly, ORC1 bound to CDC6 only beginning mid-G1 phase and into early S phase while having no interaction at all in G2-M phases, although both the proteins were found to be present in mitosis. In a reciprocal immunoprecipitation with CDC6 antibodies, ORC1 and CDC6 proteins only interacted during mid-G1 phase of the cell cycle (Figure 1E). Like Cyclin A, CDC6 protein is expressed across the cell cycle except in early G1-phase, but again our results clearly demonstrated that CDC6 was not able to co-precipitate ORC1 protein in the G2-M phase of the cell cycle. In contrast to ORC1, CDC6 immunoprecipitation did not show any interaction with ORC3 protein, but interacted with cyclin A during S-phase. The interaction between CDC6 and Cyclin A in S-phase is required for phosphorylation-mediated export of CDC6 from the nucleus to the cytoplasm to prevent re-initiation of DNA replication (Petersen et al., 1999). Thus, ORC1 and CDC6 proteins only interacted within a short window of transient interaction in mid to late G1-phase of the human cell cycle.

### CDC6 Cy motif is required for interaction with ORC1 protein

The observation that CDC6 bound to ORC1 during G1 phase but not during mitosis prompted biochemical studies to investigate their interaction. Recombinant GST-CDC6 protein expressed and purified from bacteria interacted directly with MBP-ORC1 and MBP-ORC2 proteins, but not other ORC subunits, in MBP pull-down assays (Figure 2A). This observation is consistent with structural studies of human ORC in which CDC6 was docked in between ORC1 and ORC2 through *in silico* modelling and is consistent with the yeast structure (Tocilj et al., 2017; Yuan et al., 2017). Next, many fragments of CDC6 were constructed and binding assays showed that 1-110 amino acids of CDC6 were necessary and sufficient to bind to ORC1 protein directly (Figures 2B and S2A). The data suggests that the minimal region of GST-CDC6 that bound to MBP-ORC1 protein was 1-110 (Figure S2A). Moreover, internal deletions of various small N-terminal regions (11-20, 21-30, 51-70 and 71-90) from full length GST-CDC6 protein had no effect on its binding to MBP-ORC1 protein while deletion of 90-100 and 100-110 from the full length CDC6 protein completely blocked its interaction with the ORC1 protein (Figure 2C). The ORC1 and CDC6 AAA+ ATPase regions did not interact under these conditions. Thus, the 90-110 region of CDC6 protein was critical for its interaction ORC. We noticed that the CDC6 Cy motif, CDC6^94^RRL^96^, that is required for its interaction with Cyclin E and Cyclin A was located in this region of the human CDC6 protein (Hossain and Stillman, 2016; Petersen et al., 1999; Takeda et al., 2001). We therefore tested whether it was required for binding to ORC1 and interestingly, the direct binding MBP-ORC1 protein to GST-CDC6 protein was lost in the CDC6^94^ARA^96^ (GST-CDC6 A-A) mutant when compared to wild type protein in GST pull-down assays (Figure 2D). Similarly, binding of CDC6 to Cyclin A-CDK2 was defective in this mutant protein (Figure 2E). Because the Cy motif in CDC6 is required for interaction with ORC1, we asked if the Cy motif in human ORC1 protein, which binds Cyclin A but not to Cyclin E in an ORC1^235^KRL^237^-dependent manner, was also required for ORC1 binding to CDC6 (Hemerly et al., 2009; Hossain and Stillman, 2012). However, the mutant MBP-ORC1^235^ARA^237^ (MBP-ORC1 A-A) bound to GST-CDC6 like the wild type protein, but as expected from the previous result, it did not bind to the GST-CDC6 A-A mutant protein (Figure 2F). Thus, the CDC6 Cy motif, but not the ORC1 Cy motif, was required for their direct binding, independent of any Cyclin-CDK.

**Figure 2.**
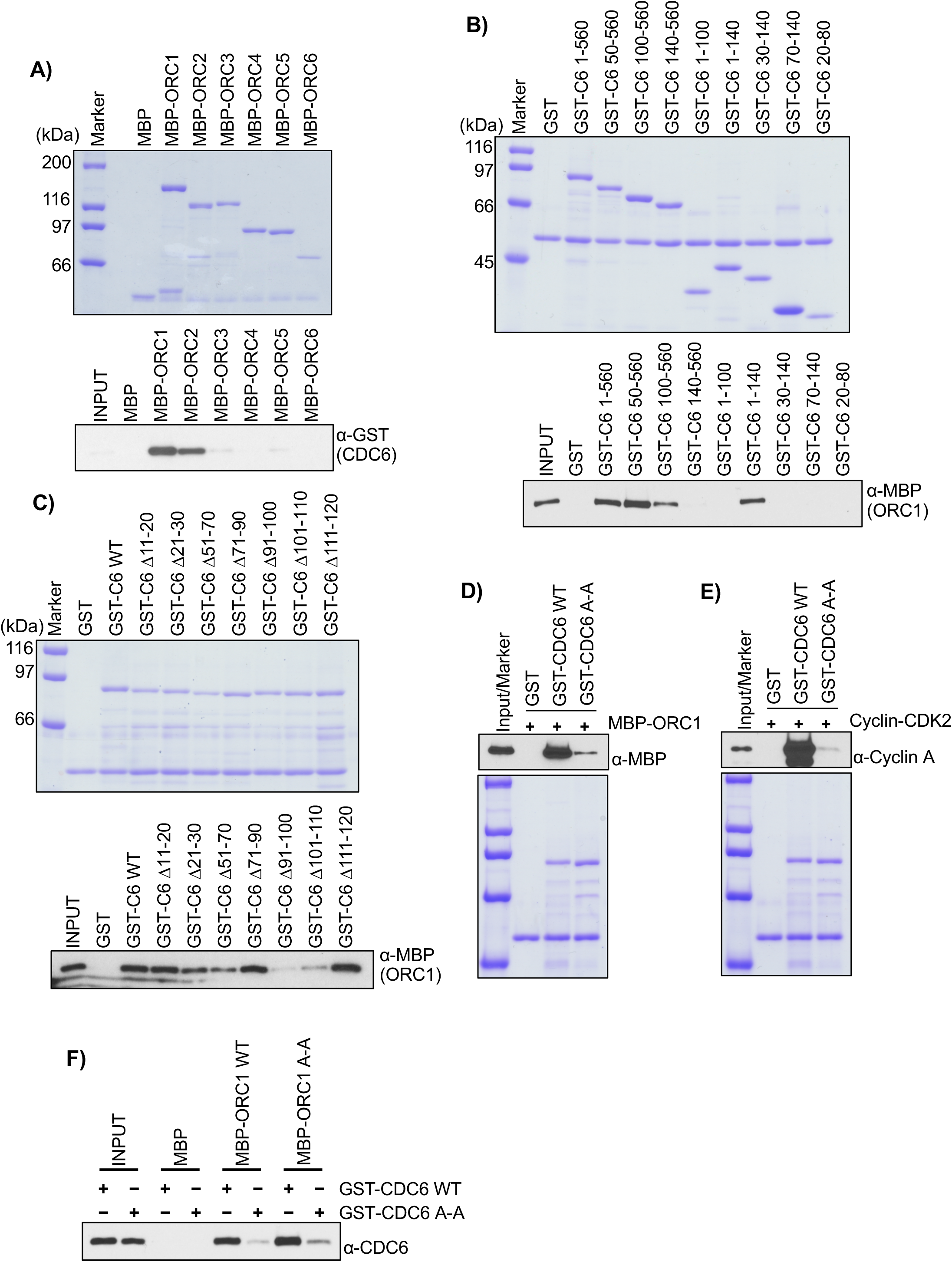
Physical interaction between ORC1 and CDC6 requires Cy motif of CDC6. (A) Purified MBP-ORC subunit proteins bound to α-MBP antibody magnetic beads were incubated with GST-CDC6 protein in an MBP pull-down assay and blotted with anti-GST antibody (bottom panel). The loading of MBP-ORC1 proteins used in pull-down assays is shown in the top panel. MBP protein served as negative control. (B-C) The recombinant GST-CDC6 full length as well as its various deletion mutant proteins bound to α-GST antibody magnetic beads were incubated with purified MBP-ORC1 protein in a GST pull-down assay and blotted with anti-MBP antibody (bottom panel). In the bottom panel of (C), equitable loading of GST-fused internal regions of CDC6 protein deleted from full length context are indicated. GST-C6 denotes GST-CDC6 protein. (D-E) The purified recombinant GST-CDC6 wild type and Cy mutant proteins bound to α-GST antibody magnetic beads were incubated with either MBP-ORC1 (D) or Cyclin A-CDK2 (E) and blotted with anti-MBP and anti-Cyclin A antibodies, respectively. The loading of GST-CDC6 proteins used in pull-down assay is shown in the top panel. GST protein served as negative control. (F) Interaction between purified recombinant full length ORC1 and CDC6 proteins in an MBP pull-down assay. MBP-ORC1 wild type or its Cy mutant protein bound to α-MBP antibody magnetic beads were incubated with either GST-CDC6 WT or Cy mutant proteins and were immunoblotted with anti-CDC6 antibody. MBP protein served as negative control.

### Cyclin A CDK2 inhibits CDC6 binding to ORC1 in mitosis, dependent on the ORC1 Cy motif and CDK phosphorylation sites

Next, the minimal regions within human ORC1 required for binding to CDC6 protein were mapped by systematically creating fragments of ORC1, expressed and purified from bacteria as MBP-tagged proteins. In MBP pull-down assays, N- and C-terminal deletions of MBP-ORC1 protein fragments indicated that CDC6 required the 100-300 region of ORC1 protein (Figure 3A and Figure 3B). Serial deletions of 20 amino acids from the C-terminal end of the 1-300 region of MBP-ORC1 demonstrated that GST-CDC6 bound to the 1-240 region of the ORC1 protein, comparable to binding of full-length ORC1 protein (Figure S3A). Upon further dissection of both N- and C-terminal deletions of the 1-300 region of MBP-ORC1 protein, GST-CDC6 protein bound to a small MBP-ORC1 fragment (165-250) while showing no interaction with either 1-200 or 250-300 regions of ORC1 protein (Figure S3B). Fine mapping of the 165-250 aa region of MBP-ORC1 protein showed that the 180-240 region of MBP-ORC1 protein was sufficient to interact directly with GST-CDC6 in an MBP pull-down assay, but not quite as well as the full length ORC1 (Figure S3C).

**Figure 3.**
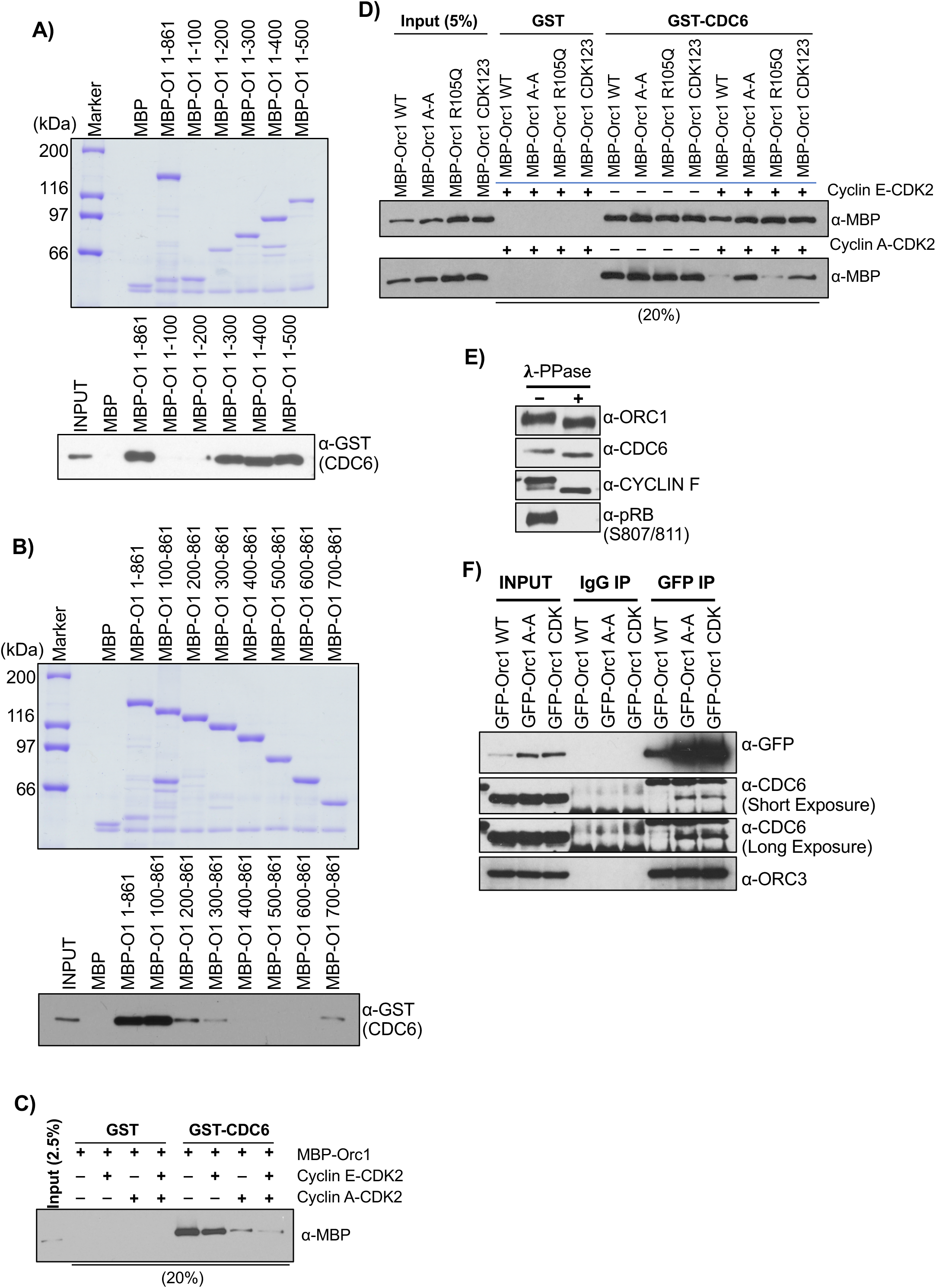
Cyclin A-CDK2 phosphorylation dependent interaction between ORC1 and CDC6 proteins. (A-B) Mapping of CDC6 interaction region in ORC1 protein. MBP-ORC1 WT or its C-terminal deletion mutant proteins (A) or N-terminal deletion mutant proteins (B) bound to α-MBP antibody magnetic beads were incubated with GST-CDC6 protein in an MBP pull-down assay and immunoblotted with anti-CDC6 antibody (bottom panel). The loading of MBP-ORC1 proteins used in pull-down assay is shown in top panel. MBP protein served as negative control. MBP-O1 denotes MBP-ORC1. (C) The GST-CDC6 proteins bound to α-GST antibody magnetic beads were incubated with MBP-ORC1 either in presence or absence of Cyclin E-CDK2 and Cyclin A-CDK2 and blotted with anti-MBP antibody in GST pull-down assay. Each of the reaction contained 1mM ATP. GST protein served as negative control. (D) In a GST pull-down assay, GST-CDC6 protein was incubated with MBP-ORC1 WT or its mutants in presence or absence of Cyclin E-CDK2 (top panel) or Cyclin A-CDK2 (bottom panel) with 1mM ATP. The western blot is probed with anti-MBP antibody and GST protein served as negative control. (E) Mitotic cells of U2OS cells are harvested after nocodazole treatment, and total cell lysates were further incubated with or without lambda protein phosphatase. The samples were used for western blotting with antibodies as indicated. (F) Mitotic cells from stable GFP-ORC1 WT or its mutant cell lines were collected after nocodazole treatment and lysed, and the cell lysates were used for immunoprecipitation with GFP antibody. The samples were further immunoblotted with CDC6 antibody (showing short and long exposure times on autorad X-ray films) and ORC3 antibody. IgG served as negative control.

The 180-240 aa region of ORC1 contains a Cy motif required for its interaction with Cyclin A in human cells, and thus we hypothesized that ORC1 and CDC6 interaction might be controlled by Cyclin A-CDK2 kinase activity. The interaction between purified ORC1 and CDC6 proteins was investigated in presence of purified G1-S kinases, Cyclin E-CDK2 and Cyclin A-CDK2. MBP-ORC1 protein was incubated with bead bound GST-CDC6 protein either in the presence or absence of purified Cyclin E-CDK2 or Cyclin A-CDK2 individually or together and the washed complexes bound to beads examined. In presence of Cyclin E-CDK2, MBP-ORC1 and GST-CDC6 proteins interacted as they did without added kinase, while their interaction was prevented by either Cyclin A-CDK2 alone or when both the kinases were present (Figure 3C). Since Cyclin A-CDK2 interacts with human ORC1 protein in a Cy motif-dependent manner, it raised the possibility that the ORC1 Cy motif might be influencing this interaction with CDC6 protein, perhaps by regulating ORC1 phosphorylation status. To better understand the role of the Cy motif in ORC1, we used an ORC1 Cy mutant (ORC1 A-A; ^235^RRL^2372^ to ^35^ARA^237^) - defective in its ability to bind Cyclin A-CDK2, an ORC1 R105Q mutant - defective in the ability of ORC1 to inhibit Cyclin E-CDK2 kinase activity, and an ORC1 CDK123 mutant (CDK1, S258A; CDK2, S273A and CDK3, T375A) - defective in phosphorylation by Cyclin E-CDK2 kinase, in GST-CDC6 pull-down assays. The interaction between either wild type or mutant MBP-ORC1 proteins with GST-CDC6 was unaffected in presence or absence of Cyclin E-CDK2 (Figure 3D, top panel). Interestingly, the binding GST-CDC6 protein to the MBP-ORC1 Cy and CDK123 mutants was not blocked by Cyclin A-CDK2, in contrast to the wild type and the R105Q mutant, implying that binding as well as phosphorylation of ORC1 by Cyclin A-CDK2 blocked its interaction with CDC6 (Figure 3D, bottom panel). Importantly, results from the MBP-ORC1 CDK123 mutant suggest that all three CDK phosphorylation sites are required by Cyclin A-CDK2 to inhibit its binding to GST-CDC6 protein as MBP-ORC1 CDK12 and MBP-ORC1 CDK3 mutant proteins behaved similar to wild type MBP-ORC1 protein (Figure S3D). Thus, we describe a previously unknown and crucial role for Cyclin A-CDK2 in regulating the interaction between human ORC1 and CDC6 proteins.

The previous immunoprecipitation results demonstrated that ORC1 and CDC6 do not interact in mitosis, although both the proteins are present. Human CDC6 is phosphorylated by Cyclin A-CDK2 (Petersen et al., 1999), as is ORC1 in Chinese Hamster cells (Li et al., 2004). To confirm whether or not human ORC1 and CDC6 proteins were phosphorylated in mitosis, U2OS cells were arrested in mitosis by treatment with various concentrations of the microtubule polymerization inhibitor nocodazole for 16 hours, and then cell extracts were prepared at different time points following release from the mitotic block for immunoblotting. Both ORC1 and CDC6 proteins migrated faster in a gel within 2 hours after nocodazole release compared to mitotic arrested cells at 0 hour, indicating post-translational modification of the proteins in mitosis (Figure S3E). Cyclin A, but not Cyclin E, is present in mitotic U2OS cells and is degraded after nocodazole release, while Cyclin E protein levels started rising at later time points (Figure S3E). Lambda phosphatase treatment of the mitotically arrested U2OS cell lysates confirmed that both human ORC1 and CDC6 were phosphorylated, since after treatment the proteins moved faster in the gel, similar to Cyclin F protein which is known to be phosphorylated in mitosis (Choudhury et al., 2017; Mavrommati et al., 2018) (Figure 3E). As a positive control for lambda phosphatase treatment, a phospho-specific RB antibody (S807/S811) was used, showing the absence of the phospho-RB protein in following phosphatase treatment.

The interaction between CDC6 in mitosis with different ORC1 mutants was tested by stably expressing tetracycline regulated GFP-ORC1 in U2OS cell lines, arresting them with nocodazole for 16 hours and preparing mitotic cell extracts. GFP immunoprecipitation from mitotic cell lysates prepared from wild type GFP-ORC1 showed no interaction with endogenous CDC6 protein, but ORC1 did bind the ORC3 protein. Interestingly, the two mutants of human ORC1 protein, GFP-ORC1 A-A (a Cy motif mutation, ORC1 ^235^A-A^237^) and GFP-ORC1 CDK123 were now able to interact with endogenous CDC6 protein along with ORC3 in GFP immunoprecipitation from their respective mitotic cell lysates (Figure 3F). Thus, we conclude that during mitosis, ORC1 binds to Cyclin A-CDK2 in an ORC1 Cy motif-dependent manner and ORC1 is phosphorylated, which prevents CDC6 from binding ORC1.

### A bi-partite region of ORC1 binds to CDC6, blocking an intramolecular interaction in the ORC1 IDR

The loss of interaction between CDC6 and ORC1 following Cyclin A-CDK2 kinase-dependent phosphorylation of ORC1 created a paradox since the minimal region of ORC1 that bound to CDC6 was 180-240 aa, but the data just described showed that phosphorylation of ORC1 outside of this region at amino acids S258; S273 and T375 prevented CDC6 from binding to ORC1 in mitosis and *in vitro* (Figures 3F and S2D). Therefore, we further investigated whether these two regions of ORC1 cooperate with each other to regulate the ORC1-CDC6 interaction. In MBP pull-down assays, wild type MBP-ORC1-180-240 and MBP-ORC1-230-400 and their corresponding mutants were incubated with GST-CDC6. Both of these partially-overlapping fragments of ORC1 bound to CDC6 protein, but the Cy mutant in the 180-240 fragment no longer bound to CDC6 protein while the same Cy mutant in the 230-400 fragment was still able to bind to CDC6 (Figure 4A). These results suggest that CDC6 binds to the 180-240 region of ORC1 in a Cy motif-dependent manner while its binding to 230-400 region is Cy motif-independent. Most interestingly, the MBP-ORC1-230-400 fragment also showed an interaction with the wild type ORC1-180-240 fragment, but not to the ORC1-180-240 Cy mutant (Figure 4B), suggesting an ORC1 intramolecular interaction. We confirmed this intramolecular ORC1 interaction by showing that MBP-ORC1-100-250 (containing 180-240) also bound to the 230-400 fragment of ORC1 in a Cy motif-dependent manner, and that Cyclin A-CDK2 kinase phosphorylation of ORC1 did not influence this binding (Figure S4A). In contrast, stoichiometric levels of CDC6 protein blocked the intramolecular interaction between the two ORC1 regions in a process that did not require any phosphorylation (Figure 4C).

**Figure 4.**
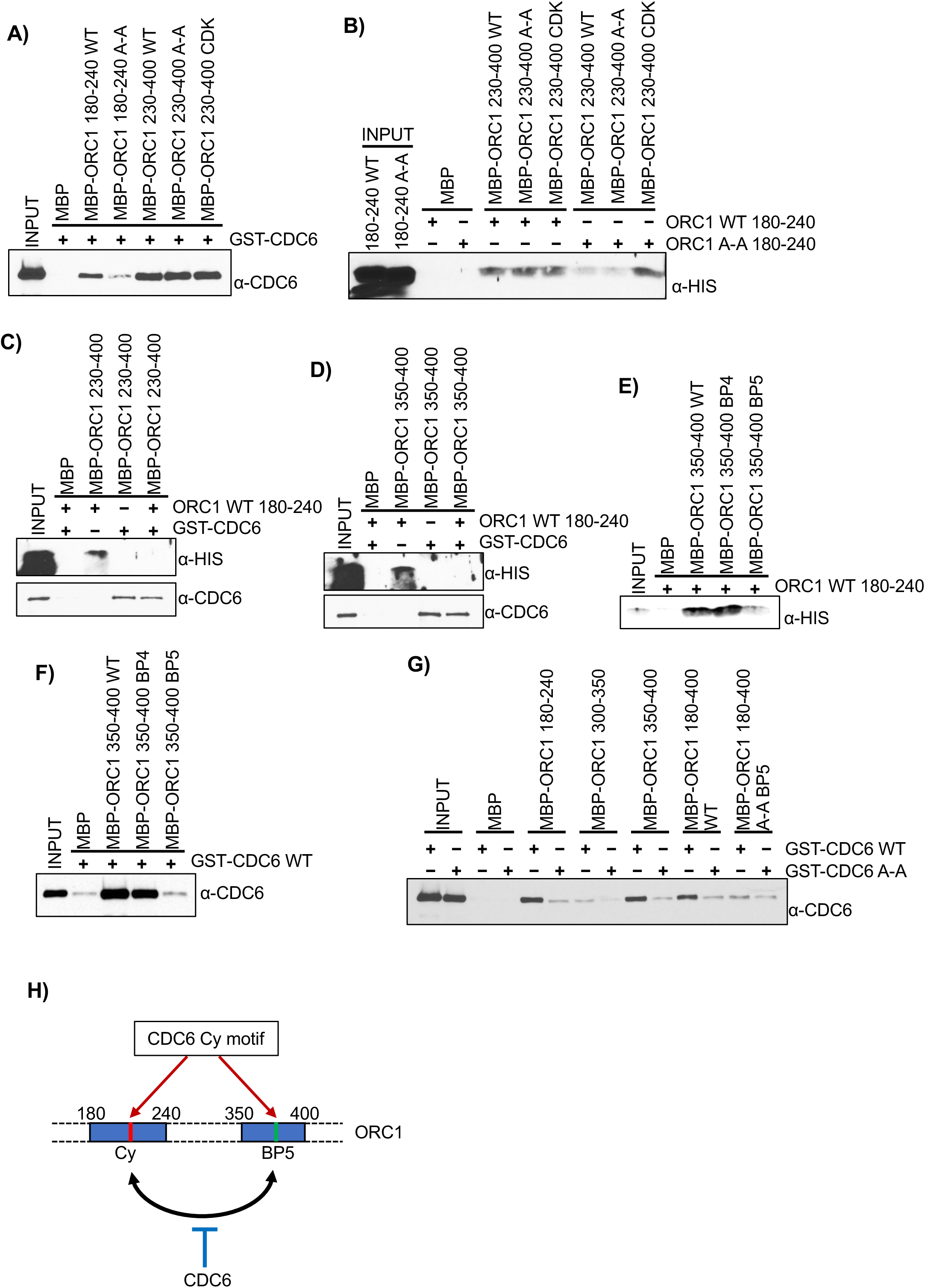
Dual mode of CDC6 binding to ORC1 protein. (A) The purified recombinant ORC1 fragments, MBP-ORC1-180-240 and MBP-ORC1-230-400 WT or its corresponding mutant proteins (A-A and CDK) bound to α-MBP antibody magnetic beads were incubated GST-CDC6 protein in MBP pull-down assays and immunoblotted with anti-CDC6 antibody. (B) MBP-ORC1-230-400 WT or its mutants bound to α-MBP antibody magnetic beads were incubated with His-tagged ORC1 protein fragments, 180-240 WT and 180-240 A-A and probed with anti-His antibody in MBP pull-down assays. (C-D) CDC6 competitively inhibits intramolecular ORC1 interaction. The recombinant ORC1 protein fragments, MBP-ORC1-230-400 (C) or MBP-ORC1-350-400 (D), bound to α-MBP antibody magnetic beads were incubated with either GST-CDC6 or ORC1-180-240-His individually or both together in MBP pull-down assays and immunoblotted with anti-CDC6 and anti-His antibodies. (E-F) The basic patch motif of ORC1 is involved in both ORC1 intramolecular-interaction and in CDC6 binding. The MBP-ORC1-350-400 WT or its basic patch motif mutant proteins (either BP4 and BP5) bound to α-MBP antibody magnetic beads were incubated with either ORC1-180-240-His (E) or GST-CDC6 protein (F) in MBP pull-down assays and the respective blots were immunoblotted with either anti-His or anti-CDC6 antibodies. (G) The various MBP-ORC1 protein fragments encompassing the 180-400 region of ORC1 bound to α-MBP antibody magnetic beads were incubated with either GST-CDC6 WT or GST-CDC6-A-A mutant proteins in MBP pull-down assays and immunoblotted with anti-CDC6 antibody. (H) Schematic showing CDC6 interacts with two distinct regions of ORC1 (180-240 and 350-400), both of which also interact with each other, and this self ORC1 interaction is competitively inhibited by wild type CDC6 protein but not its Cy (A-A) mutant. The intramolecular self-interaction within ORC1 is also lost upon ORC1-A-A mutation in the 180-240 fragment and BP5 mutation in the 350-400 fragment of the ORC1 protein.

Further mapping of the 230-400 region of MBP-ORC1 showed that the 350-400 fragment was sufficient to interact with the ORC1-180-240 in a Cy motif-dependent manner and was also competitively inhibited by GST-CDC6 protein (Figure 4D). A closer examination of the ORC1 350-400 revealed a high abundance of positively charged residues that can form basic patches involved in DNA binding (Kawakami et al., 2015) and this fragment is related to a region in *Drosophila* and *S. cerevisiae* Orc1 that is required for ORC binding to DNA (Bleichert et al., 2018; Li et al., 2018). Comparative sequence analysis shows human ORC1 contains two basic patches in this region, one previously reported as BP4 (Li et al., 2018) that is not well conserved, and another that we identified as a new basic patch and named BP5, which is highly conserved in the animal kingdom (Figure S4B). *In vitro* MBP pull-down binding assays clearly show that mutation of BP5, but not BP4, in the ORC1-350-400 fragment disrupted not only its direct interaction with ORC1-180-240 but also to GST-CDC6 (Figures 4E and 4F). Furthermore, the Cy motif in GST-CDC6 is required for binding to both the 180-240 aa and 350-400 aa regions of ORC1, since the GST-CDC6 A-A mutant lost its binding to either ORC1 fragments (Figure 4G). The 180-400 aa region of ORC1 harboring the wild type Cy motif and the MBP-ORC1-BP5 motif interacted with GST-CDC6 in a CDC6 Cy motif-dependent manner, while its interaction was lost with MBP-ORC1-180-400-A-A (Cy motif defective) and MBP-ORC1-BP5 mutant proteins in MBP pull-down assays (Figure 4G). We conclude that the Cy motif in the CDC6 protein contacts two separate regions within the ORC1 protein bearing the ORC1 Cy and BP5 motifs and that these latter two regions of ORC1 also self-interact. Thus, an ORC1 Cy motif-dependent intramolecular interaction in ORC1 is regulated by the CDC6 protein, and dependent on the CDC6 Cy motif (Figure 4H).

### Dissociation of ORC1 from CDC6 by Cyclin A-CDK2 promotes recruitment of SKP2 to ORC1 – an E3 Ubiquitin ligase for ORC1 degradation

As the levels of Cyclin A protein rise at the G1-S boundary and into S phase, the Cyclin A-CDK2 kinase disrupts ORC1 and CDC6 interaction, concomitantly leading to proteasome mediated degradation of human ORC1 protein, while phosphorylated CDC6 is exported back to cytoplasm to prevent re-initiation of DNA replication (Mendez et al., 2002; Petersen et al., 1999). It is known that human ORC1 is degraded by the SCF^SKP2^-activated proteasome complex using SKP2 as a F-box protein, but how ORC1 is targeted for degradation was not known (Mendez et al., 2002). In U2OS cells, the siRNA mediated depletion of effector molecules involved in various degradation pathways demonstrated that ORC1 degradation was solely dependent on SKP2 protein, as its loss stabilized ORC1 protein, in a similar manner to Cyclin E protein degradation, another target of the SCF^SKP2^ proteasome complex (Figure 5A). In contrast, CDC6 protein was stabilized by loss of Cdh1, which was expected (Petersen et al., 2000), but also by the loss of Cdt2 (Figure 5A) (Clijsters and Wolthuis, 2014). SETD8, which is known to be degraded by the CRL4^Cdt2^ pathway was used as a control (Centore et al., 2010; Petersen et al., 2000).

**Figure 5.**
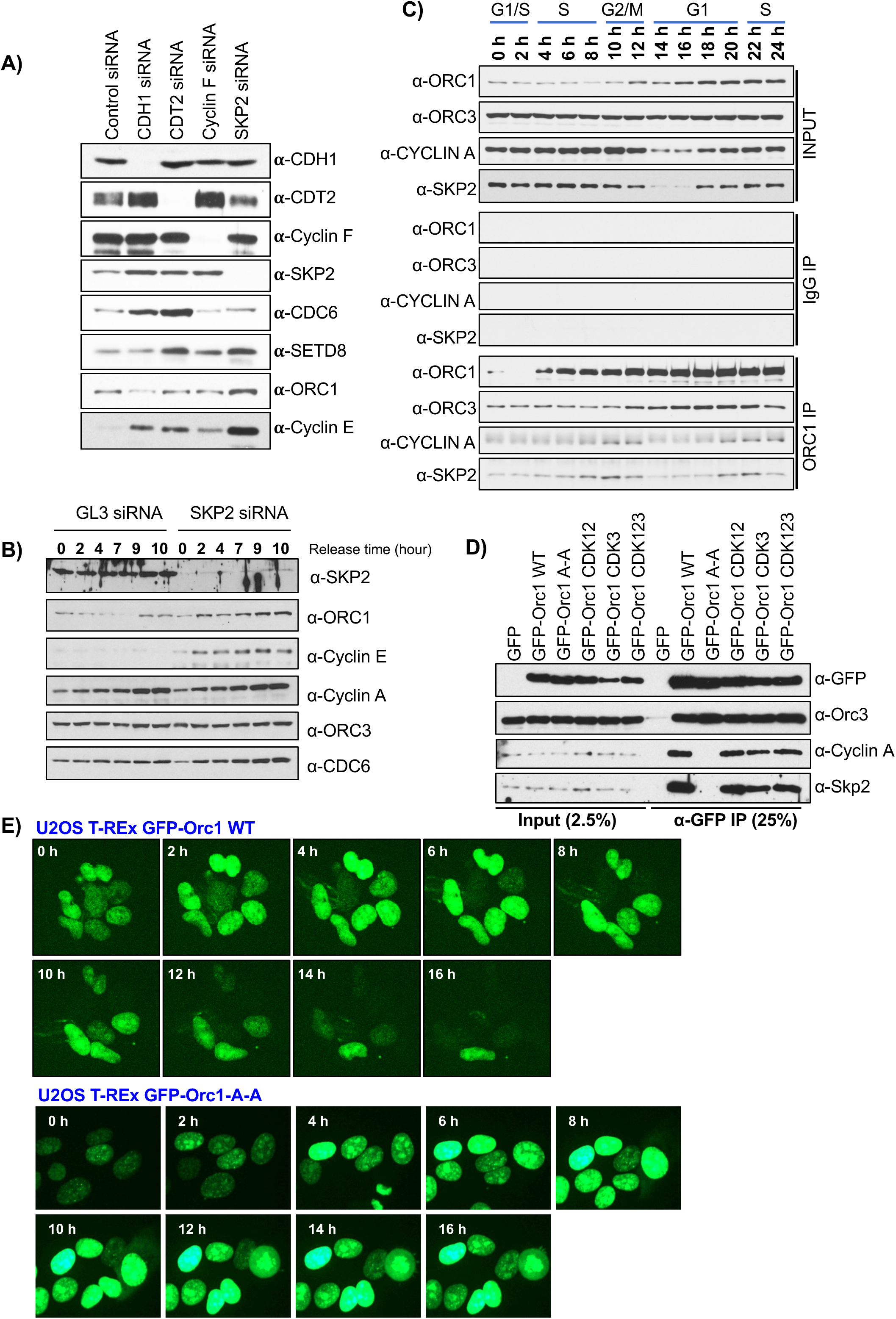
Cyclin A-CDK2 promotes ORC1 degradation in a SKP2 dependent manner. (A) The depletion of endogenous CDT2, CDH1, Cyclin F and SKP2 in asynchronous U2OS cells, following siRNA treatment for 36 hours. Total cell lysates were prepared for western blotting using antibodies as indicated. Control GL3 siRNA was used. (B) U2OS cells were transfected with either SKP2 siRNA or a control GL3 siRNA. Cells were synchronized at G1/S by double thymidine block and harvested at indicated time points after release from the block. Cell lysate were immunoblotted as indicated. (C) HeLa cells were synchronized using double thymidine block and collected at different time points after release from the block. Cells were lysed and immunoprecipitated with anti-ORC1 antibody and immunoblotted as indicated. (D) HEK293 cells were transfected with GFP-ORC1 WT or its mutants for overexpression and cells were lysed after 36 hours. Whole cell extracts were immunoprecipitated with anti-GFP antibody and immunoprecipitates were analyzed immunoblotting with indicated antibodies. (E) Tetracycline regulated GFP-ORC1 WT or its A-A mutant that were stably integrated into U2OS cell lines were used for live cell imaging for 16 hours following addition of doxycycline. The images represent time stamps from every 2-hours of movies for each cell line as indicated.

We next depleted SCF^SKP2^ in U2OS cells and synchronized them using the double thymidine block method for analysis of protein levels at various times into S phase after release from the block. ORC1 remained degraded in control siRNA treated cells, while it was stabilized following SKP2 knockdown, as also observed for Cyclin E, while the levels of Cyclin A, CDC6 and ORC3 proteins were not affected (Figure 5B). SKP2 is known to bind Cyclin A (Ji et al., 2006; Zhang et al., 1995) and using ORC1 immunoprecipitation, we found that ORC1 protein interacted with SKP2 only when Cyclin A was present across the synchronized HeLa cell cycle (Figure 5C). Cyclin A immunoprecipitation of a similar HeLa cell synchronization experiment confirmed that both ORC1 and CDC6 interacted with Cyclin A when Cyclin A and either ORC1 or CDC6 were both present in the cells (Figure S5A). Since SKP2 associated with Cyclin A, it appears that Cyclin A levels during S phase determine ORC1 protein levels.

Based on this observation, various ORC1 mutants were tested for interaction with Cyclin A and SKP2. Cells were transfected with either the GFP-ORC1 wild type plasmid, with the GFP-ORC1 A-A Cy motif mutant, or the and GFP-ORC1 CDK phosphorylation mutants and then anti-GFP antibodies were used for co-immunoprecipitation of endogenous Cyclin A and SKP2 proteins. Neither Cyclin A or SKP2 protein interacted with the GFP-ORC1 A-A Cy motif mutant protein, while mutation of the three ORC1 CDK phosphorylation sites had no effect on its binding to Cyclin A or the SKP2 protein (Figure 5D). It has been reported that the human ORC1 protein in Meier-Gorlin Syndrome (MGS) derived patient cell lines had reduced protein stability in the chromatin fraction (Bicknell et al., 2011), but we found no loss of interaction between GFP-ORC1 MGS mutants (ORC1-F89S or ORC1-R105Q or ORC1 E127E) and SKP2 protein (Figure S5B).

Since human ORC1 A-A Cy motif mutant did not bind to Cyclin A and SKP2 protein, it was possible that this mutant ORC1 protein might be resistant to SCF^SKP2^ mediated protein degradation. To test this idea, stable GFP-ORC1 wild type and mutant U2OS cell lines were constructed for live cell microscopy. The wild type GFP-ORC1 protein fluctuated during the cell cycle and was only present in mitosis and G1 phase, whereas the GFP-ORC1-A-A Cy motif mutant accumulated to high levels and was not degraded, indicating the loss of proteasome mediated control of ORC1 degradation (Figure 5E). In double thymidine synchronized and released cells that stably expressed GFP-ORC1 wild type or mutant proteins, the GFP-ORC1-A-A Cy motif mutant protein was more stable compared to wild type protein, while the levels of the GFP-ORC1 triple CDK phosphorylation were only mildly affected, confirming that GFP-ORC1-A-A mutant protein was resistant to degradation *in vivo* (Figure S5C). These results describe the discovery of a role for the Cyclin binding Cy motif of human ORC1; controlling the dynamics of ORC1 protein levels expression by binding Cyclin A and SCF^SKP2^.

### RVxF motif-dependent interaction of ORC1 with protein phosphatase PP1 controls its phosphorylation and protein stability

The observation that phosphorylated human ORC1 did not interact with CDC6 during mitosis, and the subsequent dephosphorylation of ORC1 upon mitotic exit led us to further investigate the role of protein phosphatases in this dynamic behavior. Previous reports using yeast two-hybrid systems suggest that human ORC2 interacted with protein phosphatase PP1, which in turn controlled the association ORC with chromatin by dephosphorylating ORC2 protein (Lee et al., 2014). We found that human ORC1 interacted with protein phosphatase PP1 along with its protein partner RIF1 in ORC1 immunoprecipitates from lysates of exponentially growing HeLa and U2OS cells (Figure S6A). Immunoprecipitation of all the three isoforms of GFP-PP1 (α-, β- and γ) from transiently transfected GFP-PP1 plasmids in 293 cells also detected an interaction between PP1 and endogenous ORC1 protein, along with RIF1 protein (Figure S6B) (Sreesankar et al., 2012). In MBP pull-down assays using purified proteins, GST-PP1α directly interacted with MBP-ORC1, MBP-ORC2 and MBP-CDC6, while no interaction was detected with other ORC subunits (Figure 6A). Mapping studies using N- and C-terminal deleted MBP-ORC1 proteins in MBP pull-down assays showed that the ORC1-200-300 intrinsically disordered region of ORC1 was necessary for its interaction with GST-PP1α (Figure S6C and S6D). A closer inspection of the 200-300 amino acid region of human ORC1 showed that the protein harbors a RVxF sequence (^265^KVAF^268^), a well-known PP1 binding motif, flanked by two CDK phosphorylation sites and the RVxF motif is well conserved in vertebrates (Figure 6B) (Bollen et al., 2010; Hendrickx et al., 2009). MBP-ORC1 Δ260-280, mutant protein containing an internal deletion of 20 amino acids from the full-length protein that lacks this PP1 binding motif lost its interaction with GST-PP1α protein, while deletion of 20 amino acids from adjacent regions had no effect on its binding in MBP pull-down assays (Figure S6E). Upon mutation of the wild type PP1 binding motif in MBP-ORC1 protein, from ^265^KVAF^268^ (MBP-ORC1 WT) to ^265^KAAA^268^ (MBP-ORC1-PP1^mut^), direct interaction with GST-PP1α protein was disrupted in MBP pull-down assays (Figure 6C).

**Figure 6.**
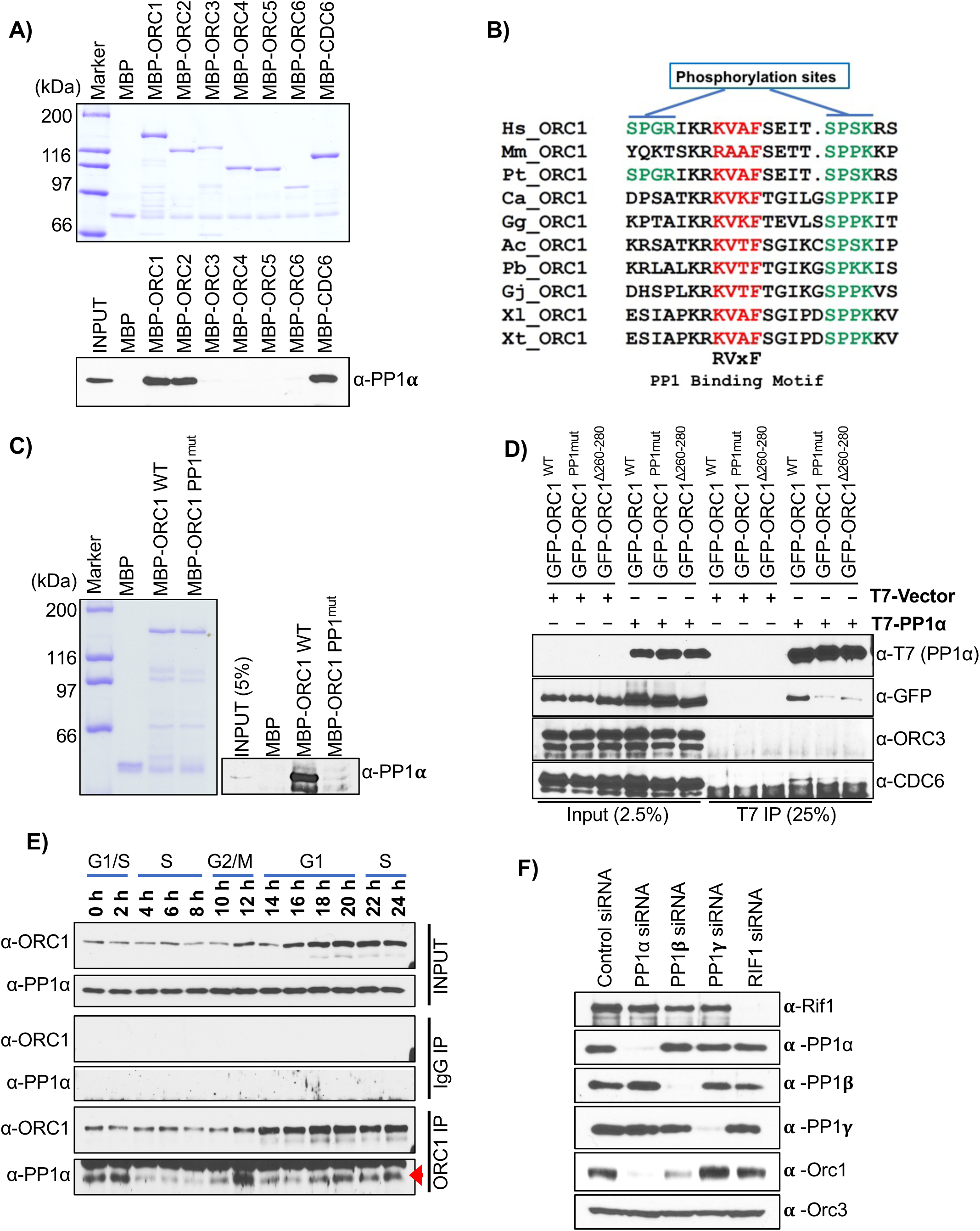
ORC1 directly binds to PP1 protein phosphatase. (A) Purified MBP-ORC subunit proteins and MBP-CDC6 protein bound to α-MBP antibody magnetic beads were incubated with purified GST-PP1α protein in an MBP pull-down assay and blotted with anti-PP1α antibody (bottom panel). The loading of MBP-fused recombinant proteins used in the pull-down assay is shown in top panel. MBP protein served as negative control. (B) Alignment of PP1 binding motif in ORC1 homologs across vertebrate species. The PP1 binding RVxF motif is highlighted in red, while the adjacent CDK phosphorylation sites are indicated in green. In multiple sequence alignment, the species representing each of the vertebrate classes are as follows: Mammals (*Homo sapiens*, Hs; *Mus musculus*, Mm; *Pan troglodytes*, Pt), Aves (*Calypte anna*, Ca; *Gallus gallus*, Gg), Reptiles (*Anolis carolinensis*, Ac; *Python bivittatus*, Pb; *Gekko japonicus*, Gj) and Amphibia (*Xenopus laevis*, Xl; *Xenopus tropicalis*, Xt). (C) MBP-ORC1 WT protein along with its corresponding PP1 binding mutant, MBP-ORC1^PP1mut^ bound to α-MBP antibody magnetic beads were incubated with purified GST-PP1α protein in MBP pull-down assay and blotted with anti-PP1α antibody (right panel). The loading of MBP-fused recombinant proteins used in the pull-down assay is shown in left panel. MBP protein served as negative control. (D) HEK293 cells were co-transfected with GFP-ORC1 WT or its PP1 binding mutants (GFP-ORC1^PP1mut^ and GFP-ORC1*^Δ^*^260-280^) with either T7-PP1α or empty T7 vector. Cell lysates from HEK293 cells transiently co-expressing the indicated constructs were immunoprecipitated with anti-T7 antibody and immunoblotted with the indicated antibodies. (E) HeLa cells were synchronized using double thymidine block and collected at different time points after release from the block. Cells harvested at indicated time points were lysed and whole cell extracts were immunoprecipitated with anti-ORC1 antibody and further immunoblotted with anti-ORC1 and anti-PP1α antibodies. Red arrow indicates the PP1 band as a non-specific band cross reacted just above the PP1 band. (F) siRNAs targeting PP1 isoforms and RIF1 were used to deplete endogenous proteins from asynchronous growing U2OS cells. Total cell lysate of siRNA treated U2OS cells were prepared after 36 hours of knockdown and immunoblotted with indicated antibodies.

We then performed a co-immunoprecipitation using HEK293 cells overexpressing T7-PP1α along with GFP-ORC1 WT or its mutants following co-transfection. The result confirmed the *in vitro* data that the RVxF motif in human ORC1 was required for its interaction with PP1α since T7-PP1α did not interact with the GFP-ORC1 Δ260-280 and GFP-ORC1 PP1^mut^ mutants compared to GFP-ORC1 wild type (Figure 6D). Furthermore, T7-PP1α immunoprecipitation did not pull-down endogenous ORC3 protein, and while its interaction with endogenous CDC6 was very weak, it too, was dependent on the RVxF motif.

The cell cycle dynamics of the interaction between ORC1 and PP1α protein was investigated by performing ORC1 immunoprecipitations from lysates of synchronized HeLa cells. ORC1 strongly interacted with PP1α during mitosis, while the interaction was slightly lowered at the G1-S transition (Figure 6E). Consistent with the observation that ORC1 protein was phosphorylated in mitosis, *in vitro* phosphorylation of MBP-ORC1 WT or its A-A Cy motif and CDK phosphorylation mutants by either Cyclin E-CDK2 or Cyclin A-CDK2 kinases did not impair its binding to GST-PP1α in MBP pull-down assays, while MBP-ORC1 PP1^mut^ with GST-PP1α did not bind even after phosphorylation (Figure S6F). Thus, our data demonstrates that human ORC1 phosphorylation does not affect its interaction with PP1α during mitosis.

Based on the results above, we treated U2OS cells with the protein synthesis inhibitor, cycloheximide, for different durations and compared the protein stability of ORC1 and CDC6 proteins. CDC6 protein was rapidly degraded, similar to Cyclin F, but the turn-over rate for ORC1 and Cyclin E proteins was slower (Figure S6G). Furthermore, to investigate the role of protein phosphorylation in human ORC1 protein degradation, cells were treated with roscovitine, a CDK inhibitor. Upon treatment, U2OS cells showed a rapid decrease in the stability of CDC6 protein with time, leading to the inference that the phosphorylation of endogenous CDC6 stabilizes the protein (Mailand and Diffley, 2005), as previously also observed in case of Cyclin F protein. However, roscovitine treatment of U2OS cells showed an opposite effect on endogenous ORC1 protein, with a concomitant increase in the protein levels, indicating that phosphorylation of human ORC1 promoted its degradation, while no effect was seen on Cyclin E protein levels (Figure S6H).

Depletion of individual PP1 isoforms or RIF1 using siRNAs in U2OS cells showed that upon knockdown of PP1α the endogenous ORC1 protein level was reduced, and depletion of PP1β reduced ORC1 to a lesser extent and reducing PP1γ had no effect (Figure 6F). The results indicated that PP1α controls phosphorylation mediated degradation of ORC1 in mammalian cells. We did not observe any effect on ORC1 protein levels upon RIF1 depletion in contrast to a previous report indicating RIF1 mediated recruitment of RIF1-PP1 complex in controlling ORC1 protein stability (Hiraga et al., 2017).

### Dominant negative phenotype of human ORC1 mutants defective in binding Cyclin A-CDK2 kinase and PP1 protein phosphatase

The phenotypes associated with mutants in the ORC1 Cy motif, the three CDK sites and the PP1 binding site were investigated. Cell proliferation and cell cycle profiles of stable U2OS cell lines harboring tetracycline inducible GFP-ORC1 WT or mutants were monitored for 72 hours after depletion of endogenous ORC1 protein using a 3’-UTR specific ORC1 siRNA. Compared to control siRNA, the depletion of endogenous ORC1 from U2OS cells significantly retarded its proliferation, while in the GFP-ORC1 WT U2OS cell line, GFP-ORC1 rescued the proliferation defect in siORC1 treated cells (Figures 7A & 7B). In contrast, expression of either GFP-ORC1 mutants - GFP-ORC1 A-A, GFP-ORC1 CDK123 or GFP-ORC1^PP1mut^ were not able to rescue the proliferation defect associated with endogenous ORC1 protein depletion (Figure 7A). Interestingly, when the GFP-ORC1 mutants were expressed, the cells proliferated slower in both siORC1 treated and even control siRNA treated cells, demonstrating a dominant negative phenotype associated with these mutant proteins. Flow cytometry data shows that after doxycycline induction, the GFP-ORC1 A-A and GFP-ORC1 CDK123 cell lines had constant and high GFP expression of these mutant proteins in S phase, while the expression of GFP-ORC1 WT and GFP-ORC1^PP1mut^ remained comparable and very low (Figure 7C and Figure S7). During the G2/M phase, the expression of GFP-ORC1 A-A and GFP-ORC1 CDK123 mutants in the induced cell lines significantly increased compared to GFP-ORC1 WT, while the GFP-ORC1^PP1mut^ was only marginally elevated (Figure 7D and Figure S7).

**Figure 7.**
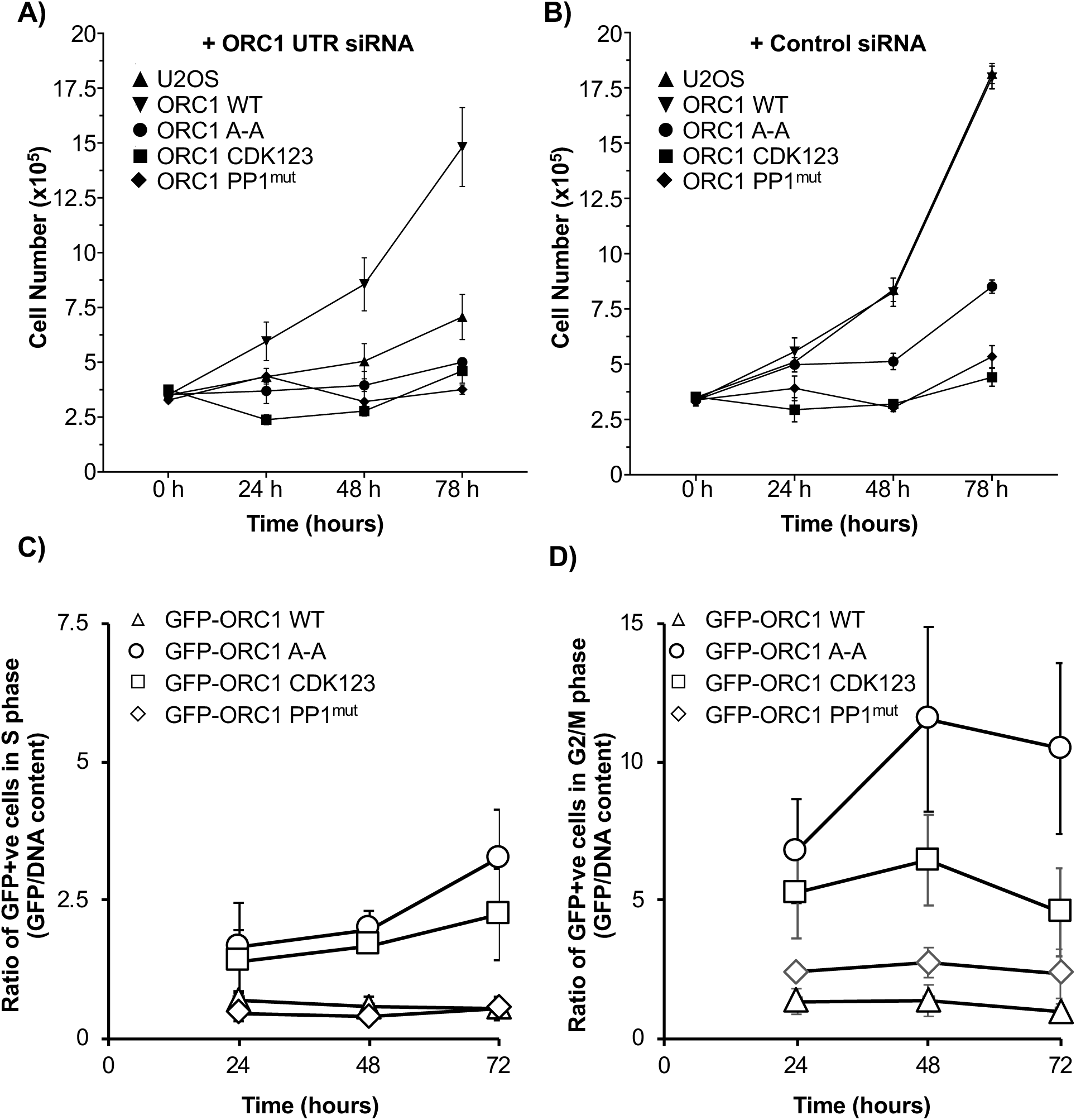
Reduced cell proliferation of human ORC1 mutants. (A-B) Doxycycline induced U2OS stable cell lines expressing GFP-ORC1 WT and its mutants (GFP-ORC1 A-A, GFP-ORC1 CDK123 and GFP-ORC1 PP1^mut^) along with control U2OS cells were treated with 3’-UTR specific ORC1 siRNA to deplete endogenous ORC1 (A) or with control siRNA (B). The total number of viable cells for each cell line are counted every 24 hours for 72 hours. Error bars represent SD from three independent experiments. (C-D) The mean fluorescence intensity of GFP in U2OS stable cell lines expressing GFP-ORC1 WT and its mutants (GFP-ORC1 A-A, GFP-ORC1 CDK123 and GFP-ORC1 PP1^mut^) from S phase (C) and G2/M phase (D) were analyzed with flow cytometry and normalized it to the DNA content of respective phases of the cell cycle. The y-axis represents GFP intensity normalized to DNA content for 72 hours at every 24-hour time interval. Error bars represent SD from three independent experiments. The original 2-D flow cytometry from one experiment is in Figure S7.

## Discussion

The ORC1 and CDC6 IDRs contain multiple SLiMs, including in ORC1 the nuclear localization signal, the PP1 binding site, the Cy motif and the BP5 motif that promotes ORC1 intramolecular interaction and binding to CDC6 (Figure S8). The CDC6 IDR contains a multifunctional Cy motif and a degron (Figure S8).

The Cyclin binding Cy motif has been long recognized as a short linear peptide motif (SLiM) within both kinase substrate and inhibitor proteins, that interacts with the many isoforms of Cyclin-CDKs (Adams et al., 1996; Petersen et al., 1999; Schulman et al., 1998; Takeda et al., 2001; Wohlschlegel et al., 2001; Wood and Endicott, 2018). Investigation of when and how ORC and CDC6 interact during the cell division cycle in human cells led us to uncover broader roles for Cy motifs that are independent of Cyclin binding protein partners in some instances. Moreover, the Cy motifs in ORC1 and CDC6 have different specificities and their function varies with the stage in the cell division cycle. Starting in mitosis, where both ORC1 and CDC6 are present, these proteins cannot interact because of Cyclin A-CDK2 mediated phosphorylation at three kinase target residues in ORC1 (S258A; S273A and T375A). The inhibition of ORC1-CDC6 interaction requires the Cy motif in ORC1, but not the Cy motif in CDC6, suggesting that in mitosis, the ORC1 Cy motif functions in its classic Cyclin binding mode, recruiting the Cyclin-CDK kinase and accelerating the phosphorylation of ORC1. Both the Cy motif and the three CDK phosphorylation sites are in the IDR of human ORC1 protein (Figure S8), a common feature of SLiMs (Van Roey and Davey, 2015; Van Roey et al., 2014). This function of the ORC1 Cy motif in mitosis is akin to the binding of Cy motifs in the CDK inhibitor proteins p21^CIP1^ and p27^KIP1^ or in the substrate CDC6 which bind to a hydrophobic patch containing the consensus MRAIL amino acid sequence that is highly conserved among many Cyclin proteins, either inhibiting the kinase or accelerating phosphorylation of the substrate protein (Furstenthal et al., 2001; Russo et al., 1996; Schulman et al., 1998; Wood and Endicott, 2018). We suggest that the inability of ORC1 to bind CDC6 in mitosis contributes to the prevention of premature assembly of the pre-RC, similar to the inhibition of CDT1 by Geminin during mitosis (Lygerou and Nurse, 2000; Wohlschlegel et al., 2000). However, mutation of this sequence in ORC1 has multiple phenotypes, depending on the cell cycle context, implicating the diverse roles of the ORC1 Cy motif.

Upon transition to the next G1 phase, human ORC1 is inherited into the two daughter cells by virtue of its binding to the mitotic chromosomes, and only then does it bind to the other ORC subunits (Kara et al., 2015; Siddiqui and Stillman, 2007). The mechanism of assembly of ORC proteins in human cells is different from that observed in Chinese hamster cells, since it has been reported that hamster CgOrc1 is stable throughout the cell cycle, is ubiquitylated upon G1/S phase transition, but is not degraded (Li et al., 2004; McNairn et al., 2005; Okuno et al., 2001). Moreover, the current understanding of binding of Orc1 in Chinese hamster CHO cells is confusing, with one report stating the Cyclin A-CDK2 blocks Orc1 from binding to chromatin during mitosis while another reporting that multiple CHO ORC subunits, including Orc1, bind to mitotic chromosomes (Li et al., 2004; Okuno et al., 2001).

The observation that in human cells ORC1 only binds to CDC6 in mid to late G1 phase and that this interaction requires the CDC6 Cy motif suggests a new role for the Cy motif, as a ligand for Cyclin-independent protein-protein interactions. CDC6 bound to two separate regions in the IDR of ORC1, the first region (180-240) contains the ORC1 Cy motif and the other has the conserved BP5 region (350-400). The CDC6 Cy motif, which is itself located in the protein’s IDR (Figure S8), is required for binding to both the regions of ORC1. Interestingly, the two domains of ORC1, 180-240 and the BP5 region bind to each other in an intramolecular interaction that is disrupted in the presence of CDC6. It is likely that CDC6 binds in concert with Cyclin E-CDK2 since we have shown that both bind to ORC1 *in vitro*, forming a ternary complex (Hossain and Stillman, 2016). Precisely at the time when ORC1 binds to CDC6, the MCM2-7 proteins are loaded onto chromatin in human cells (Mendez and Stillman, 2000; Mendez et al., 2002). Thus, CDC6 and Cyclin E synthesis and hence pre-RC formation in human cells occurs only after cells have committed to re-enter the cell division cycle, a decision regulated by activation of Cyclin D-CDK4/6 (Johnson and Skotheim, 2013; Narasimha et al., 2014) and also involving ORC1 binding to the Retinoblastoma protein and the histone methyltransferase SUV39H1 (Hossain and Stillman, 2016). These results are consistent with the earlier demonstration that overexpression of CDC6 and Cyclin E shortens G1 phase and accelerates the licensing of DNA replication (Coverley et al., 2002). However, over-expression of Cyclin E causes replication stress and chromosome segregation errors leading to genome instability and cancer (Ekholm-Reed et al., 2004; Macheret and Halazonetis, 2018; Petropoulos et al., 2019; Teixeira and Reed, 2017; Teixeira et al., 2015).

The finding that the Cy motif in CDC6 is required for binding to ORC1 in an interaction that does not involve Cyclin proteins expands the role of the Cy motif as a protein interaction motif. The CDC6 Cy motif lies within a predicted IDR and we are currently investigating its structure as it binds the two domains of ORC1. This role for the Cy motif acting as a protein interaction motif or ligand is analogous to the SRC Homology 3 Domain (SH3) acting as a ligand for proline rich sequences in adaptor proteins during signal transduction or the LxCxE motif acting as a ligand for binding to the retinoblastoma protein (Van Roey and Davey, 2015; Van Roey et al., 2014). We suggest that Cy motifs in other Cyclin-CDK substrates may mediate Cyclin-independent protein-protein interactions similar to that found in the ORC1-CDC6 interaction.

It is particularly noteworthy that the ORC1 BP5 region (aa 350-400) that intramolecularly binds to the ORC1 180-240 region, both of which are present in the ORC1 IDR (Figure S8), is related in sequence to a short region of the *S. cerevisiae* Orc1 that binds to DNA when ORC first encounters origin DNA (Li et al., 2018). This protein-DNA interaction is lost when yeast ORC binds to Cdc6 and assembles into an ORC-Cdc6-Cdt1-Mcm2-7 (OCCM) pre-RC loading intermediate (Yuan et al., 2017), indicating its dynamic DNA binding nature. The IDR of *Drosophila* ORC1 has also been shown to be required for ORC binding to DNA and in the presence of DNA it forms a phase transition condensate (Bleichert et al., 2018; Matthew W Parker, 2019). We suggest that the Cy motif in CDC6, in cooperation with Cyclin E-CDK2 controls these ORC1 protein-DNA and protein-protein interactions and phase transitions during pre-RC assembly in G1 phase. Moreover, we suggest that the multiple SLiMs in the ORC1 and CDC6 IDRs bind and bring together the AAA+ ATPase domains of these proteins that then form an intimate, ATP-dependent interaction for pre-RC assembly of MCM2-7 helicase subunits.

Following pre-RC assembly at origins, as Cyclin A-CDK2 is resynthesized at the G1/S phase transition to activate DNA replication, we observed that Cyclin A-CDK2 bound to ORC1 via its Cy motif and co-recruited SKP2 to the complex. SKP2 depletion increases ORC1 levels, consistent with our previous studies on cell cycle regulated ORC1 stability in human cells (Mendez et al., 2002). The binding of SKP2 and Cyclin A-CDK2 to ORC1 is not affected by the three CDK sties that influence its interaction with CDC6 in mitosis, suggesting that the Cy motif functions in a different manner at the G1/S phase transition than it does in mitosis. Furthermore, in early G1 phase, the APC^CDH1^ ubiquitin ligase antagonizes the SKP2 ubiquitin ligase (Bashir et al., 2004; Wei et al., 2004), thereby preventing premature ORC1 degradation. Thus, the ORC1 Cy motif, at the G1/S phase transition functions as a trigger for ORC1 destruction and is thus part of the ORC1 degron. This observation uncovers a third role for a Cy motif, as a direct regulator of protein degradation, analogous to other small linear peptide motifs, the APC/C degron box RxxLxxΦ, or the PIP (PCNA interacting protein) degron that mediates recruitment of the CRL4^Cdt2^ E3 ubiquitin ligase (Van Roey and Davey, 2015). It is striking that the CDC6 degron for APC/C mediated degradation is related in sequence to the ORC1 Cy motif degron that is targeted by SCF^SKP2^ (Figure S8). We suggest that these two mechanisms have a common evolutionary origin.

In addition to the Cy motifs in ORC1 and CDC6 and the BP5 region in ORC1 that are located in the IDRs of these proteins, we found another SLiM (^265^KVAF^268^) within the IDR of ORC1 that conforms to the consensus RVxF binding motif found in other PP1 substrates (Figure S8). Mutation of this motif prevented PP1 from binding to ORC1 and removal of phosphate groups from ORC1 in early G1 phase was partially impaired. Furthermore, the depletion of protein phosphatases from cells also led to degradation of endogenous human ORC1 protein implicating the role of ORC1 protein phosphorylation in promoting its stability or chromatin association. These results show that human ORC1 can directly interact with PP1 and that this interaction is not dependent on RIF1 as has been previously reported (Hiraga et al., 2017). Expression of mutant ORC1 that cannot bind PP1 in human cells caused a dominant inhibition of cell proliferation, suggesting that removal of the phosphate groups on ORC1 is critical for its CDC6 binding activity in mid G1 phase of cell cycle as well as its degradation in S-phase.

Multiple SLiMs are often found in IDRs, including IDRs that regulate protein kinases and many protein-protein assemblies such as transcription, splicing and protein location in cells (Gogl et al., 2019; van der Lee et al., 2014) and this is also the case for the two ATPase subunits of the human replication initiator ORC-CDC6. The diverse functions of the ORC-CDC6 SLiMs, as protein-protein interaction motifs, enzyme accelerators for both kinases and a phosphatase, as a degron and as DNA binding motifs underscore the context dependency of these modules and show a remarkable complexity of function within the IDR. We suspect that the IDRs of ORC1 and CDC6 harbor other SLiM activities.

## Acknowledgements

We thank Patricia Wendel and Ariana Amendolara for technical help and Leemor Joshua-Tor for discussions and comments on the manuscript. The antibody and the microscopy shared resources of the Cold Spring Harbor Laboratory Cancer Center (P30CA045508). This research was supported by grants from the National Cancer Institute (PO1CA113106) and the National Institute of General Medical Sciences (RO1GM45436).

## Author Contributions

M. H., K. B. and B. S. conceived and designed the experiments and analyzed the data. M. H. and K. B. performed the experiments. M. H., K. B. and B. S. wrote and edited the paper, as well as designing and drawing the figures.

## Supplemental information titles and figure legends

**Figure S1.**
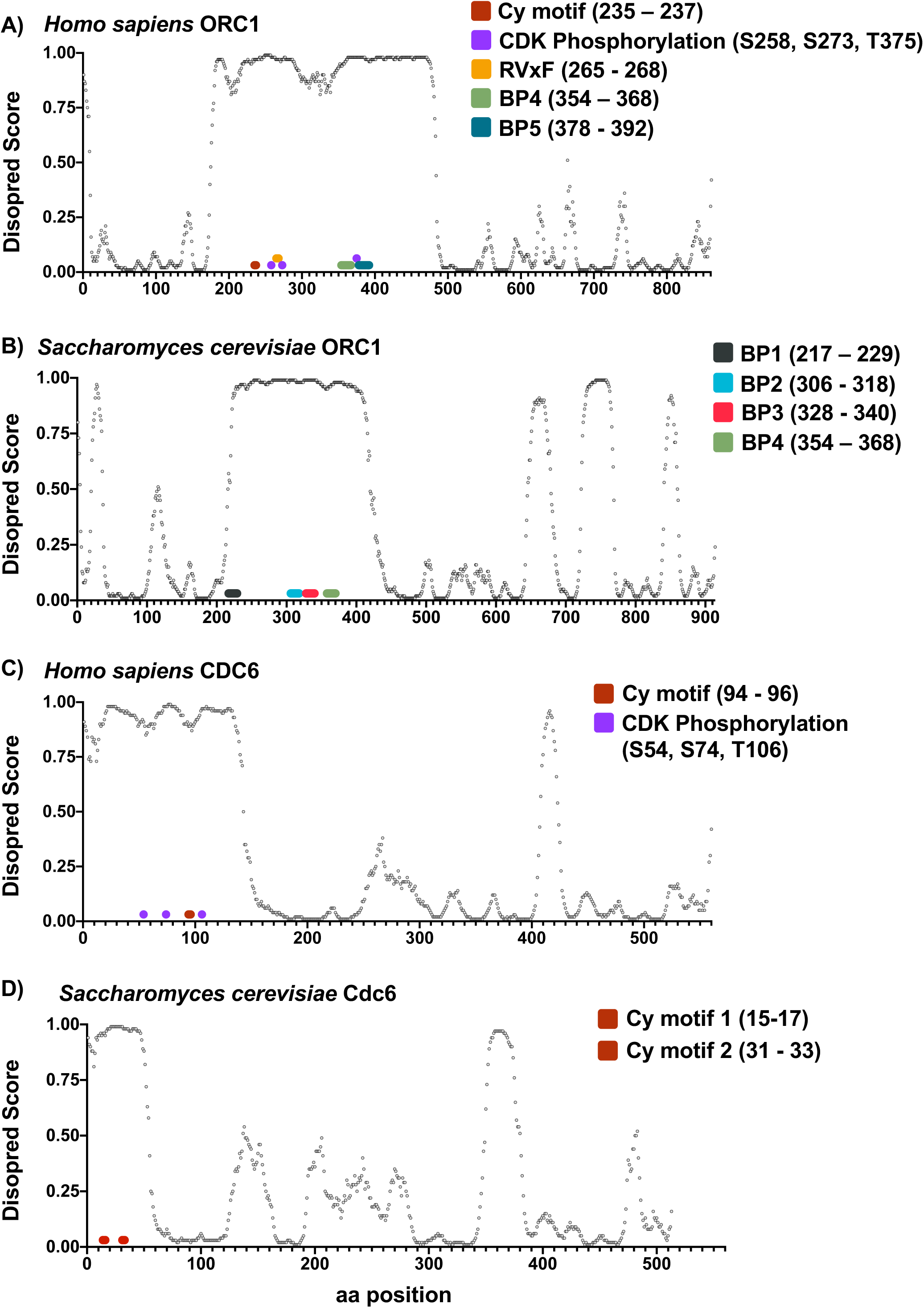
Predicted intrinsically disordered regions of ORC1 and CDC6 harbor Cy motifs. Human and Yeast reference sequences of ORC1 and CDC6 (A) human ORC1 - NP_004144.2; (B) yeast ORC1 - NP_013646.1; (C) human CDC6 - NP_001245.1 and (D) yeast CDC6 - NP_012341.1 were put through a disorder prediction algorithm, DISOPRED3 hosted on the PSIPRED server (http://bioinf.cs.ucl.ac.uk/psipred/; Jones, D.T., and Cozzetto, D. (2015). DISOPRED3: precise disordered region predictions with annotated protein-binding activity. Bioinformatics *31*, 857–863). Intrinsic disorder score for each amino acid position is shown as scatter plots. Motif positions are shown as indicated in the individual legends.

**Figure S2.**
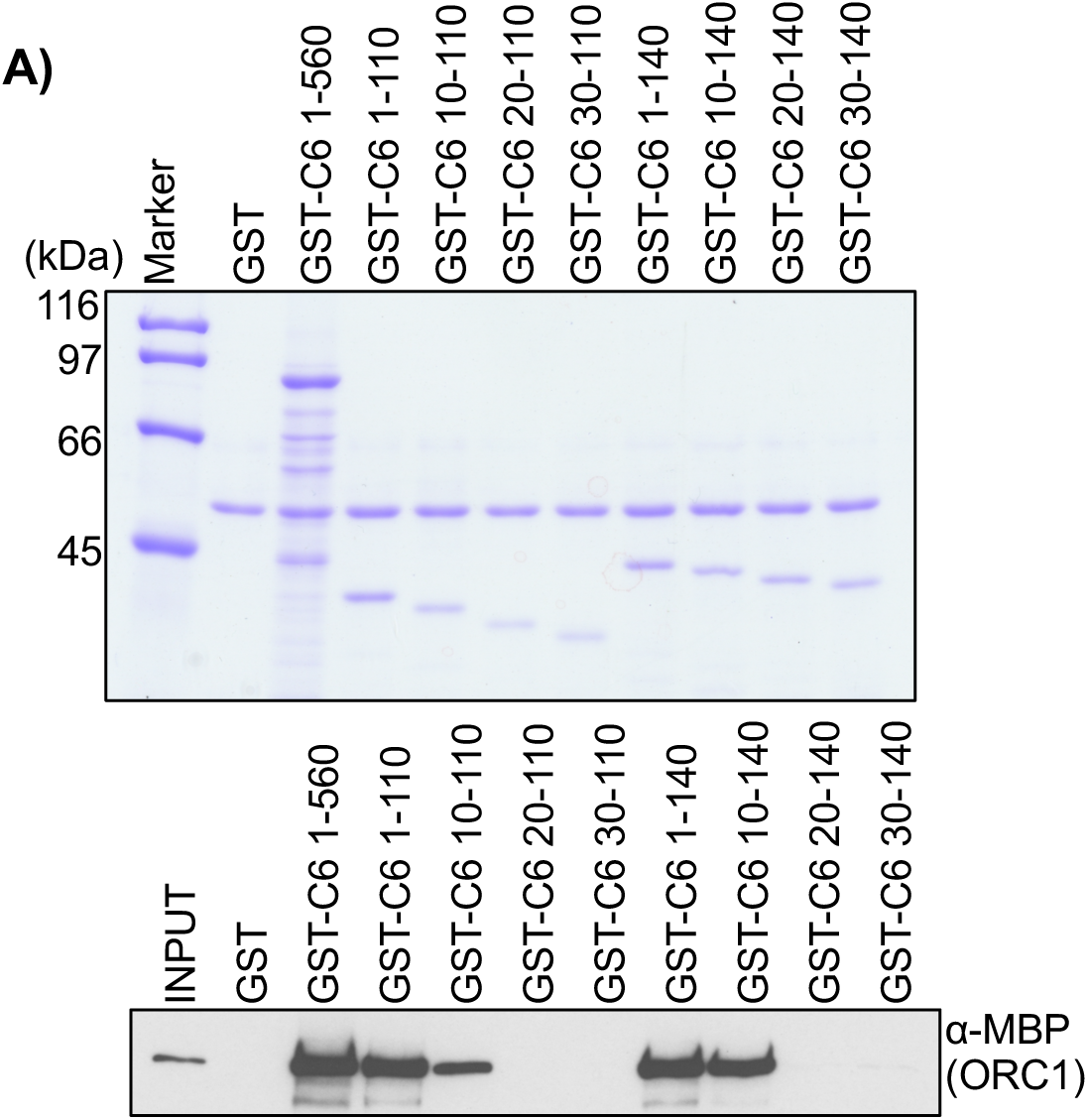
ORC1 binds the N-terminus of CDC6 protein, Related to Figure 2. (A) Purified GST-CDC6 WT (1-560 aa) and its N-terminal deletion mutant proteins as indicated, bound to α-GST antibody magnetic beads were incubated with purified MBP-ORC1 protein in GST pull-down assays and blotted with anti-MBP antibody (bottom panel). Equitable GST-fusion proteins used in pull-down assays are shown in the top panel. GST served as negative control.

**Figure S3.**
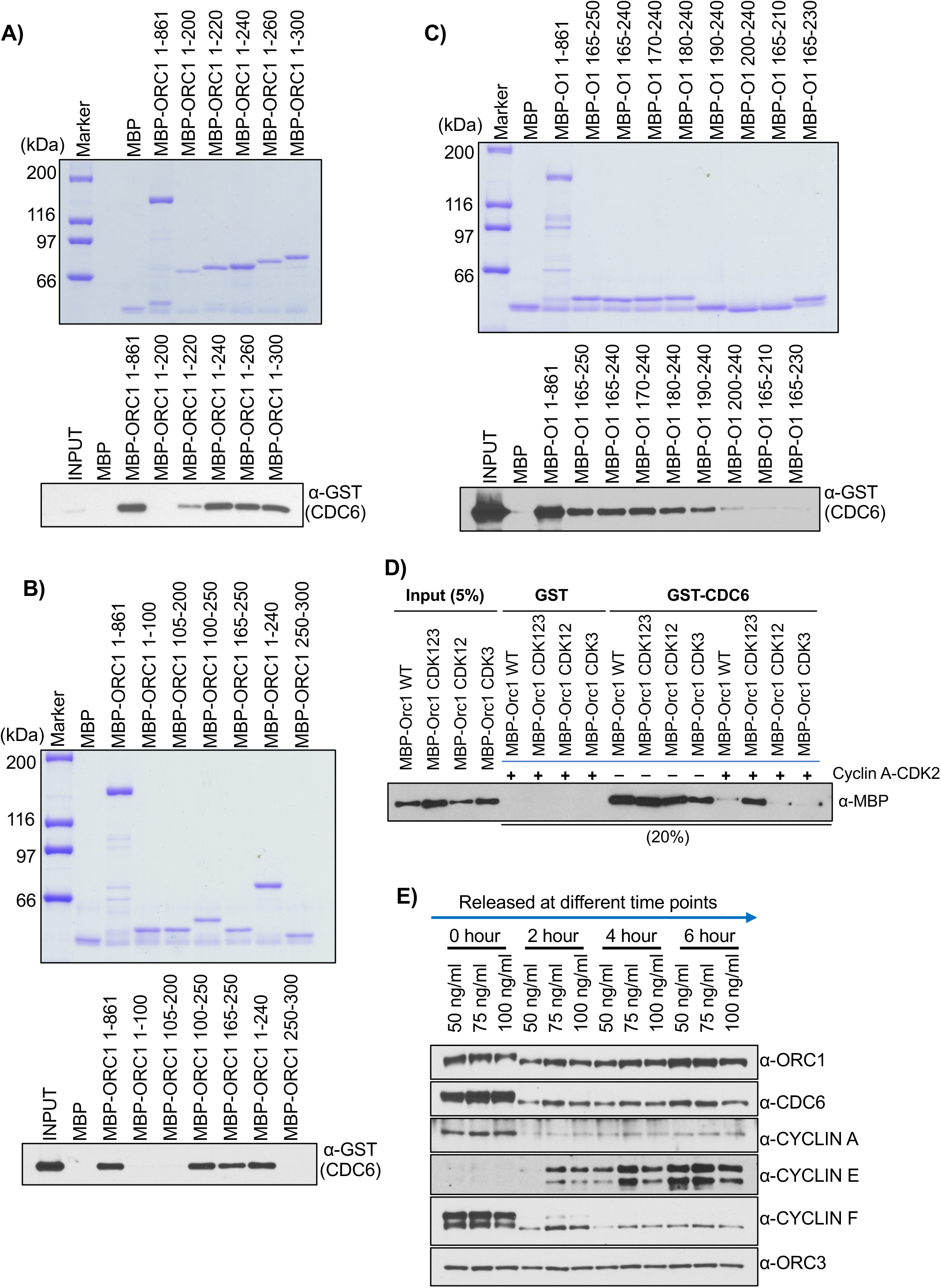
Mapping of ORC1 region binding to CDC6 and ORC1-CDC6 interaction is regulated by Cyclin A-CDK2 phosphorylation, Related to Figure 3. (A-C) Mapping of CDC6 interaction region in ORC1 protein. MBP-ORC1 WT or its various deletion mutant proteins as indicated, bound to α-MBP antibody magnetic beads were further incubated with purified GST-CDC6 protein in MBP pull-down assays and immunoblotted with anti-CDC6 antibody (bottom panels). MBP-ORC1 proteins used in pull-down assays are shown in the top panels. MBP protein served as negative control. (D) In a GST pull-down assay, recombinant GST-CDC6 proteins bound to α-GST antibody magnetic beads were incubated with purified MBP-ORC1 WT or its CDK mutant proteins in presence or absence of Cyclin A-CDK2, in presence of 1mM ATP. The western blot is probed with anti-MBP antibody and GST protein served as negative control. (E) U2OS cells were synchronized in mitosis with different concentrations of Nocodazole as indicated, isolated mitotic cells were released and harvested at different time points into the cell cycle and total cell lysates were analyzed by immunoblotting with different antibodies as indicated.

**Figure S4.**
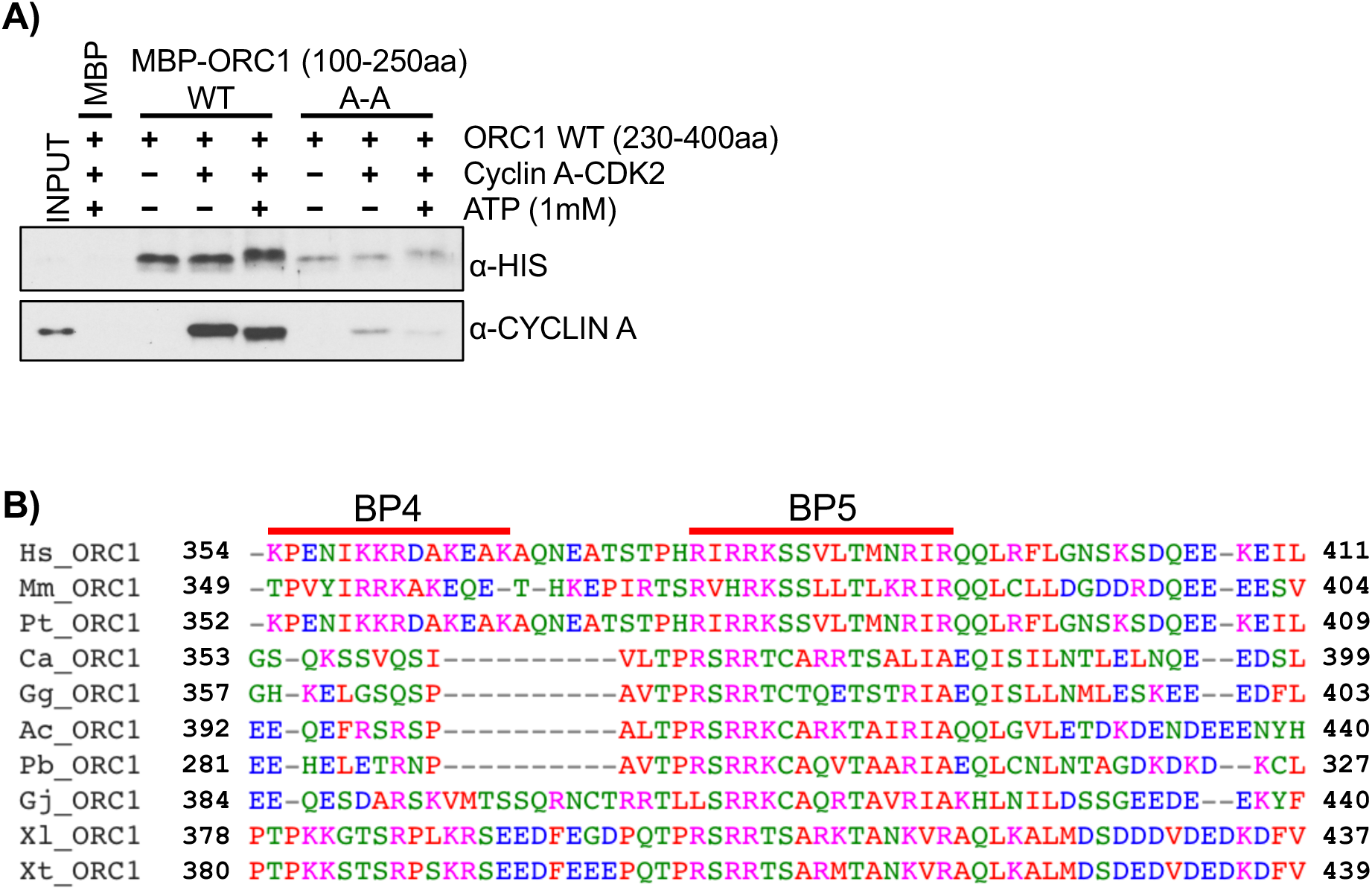
Intramolecular interaction of ORC1 protein, Related to Figure 4. (A) ORC1 fragments, MBP-ORC1 100-250 aa WT and MBP-ORC1 100-250 aa A-A mutant proteins bound to α-MBP antibody magnetic beads were incubated with ORC1 230-400-His protein either alone or in presence of Cyclin A-CDK2 with or without 1mM ATP in MBP pull-down assays. The pull-down assays were western blotted with anti-HIS and anti-CDC6 antibodies. (B) Different fragments of ORC1 protein from 230-400aa region were used in MBP pull-down assay. The MBP-ORC1 protein fragments as indicated bound to α-MBP antibody magnetic beads were incubated with either ORC1 180-240-His WT or ORC1 180-240-His A-A mutant proteins and immunoblotted with anti-His antibody. (C) Multiple sequence alignment of basic patch motifs within the 350-400 region of human ORC1 protein and its corresponding homologs across vertebrate species. The two basic motifs (BP4 and BP5) in human ORC1 protein are indicated by red lines above the alignment. Conserved basic residues in the motifs are colored in pink. In multiple sequence alignment, the species representing each of the vertebrate classes are abbreviated as follows: Mammals (*Homo sapiens*, Hs; *Mus musculus*, Mm; *Pan troglodytes*, Pt), Aves (*Calypte anna*, Ca; *Gallus gallus*, Gg), Reptiles (*Anolis carolinensis*, Ac; *Python bivittatus*, Pb; *Gekko japonicus*, Gj) and Amphibia (*Xenopus laevis*, Xl; *Xenopus tropicalis*, Xt).

**Figure S5.**
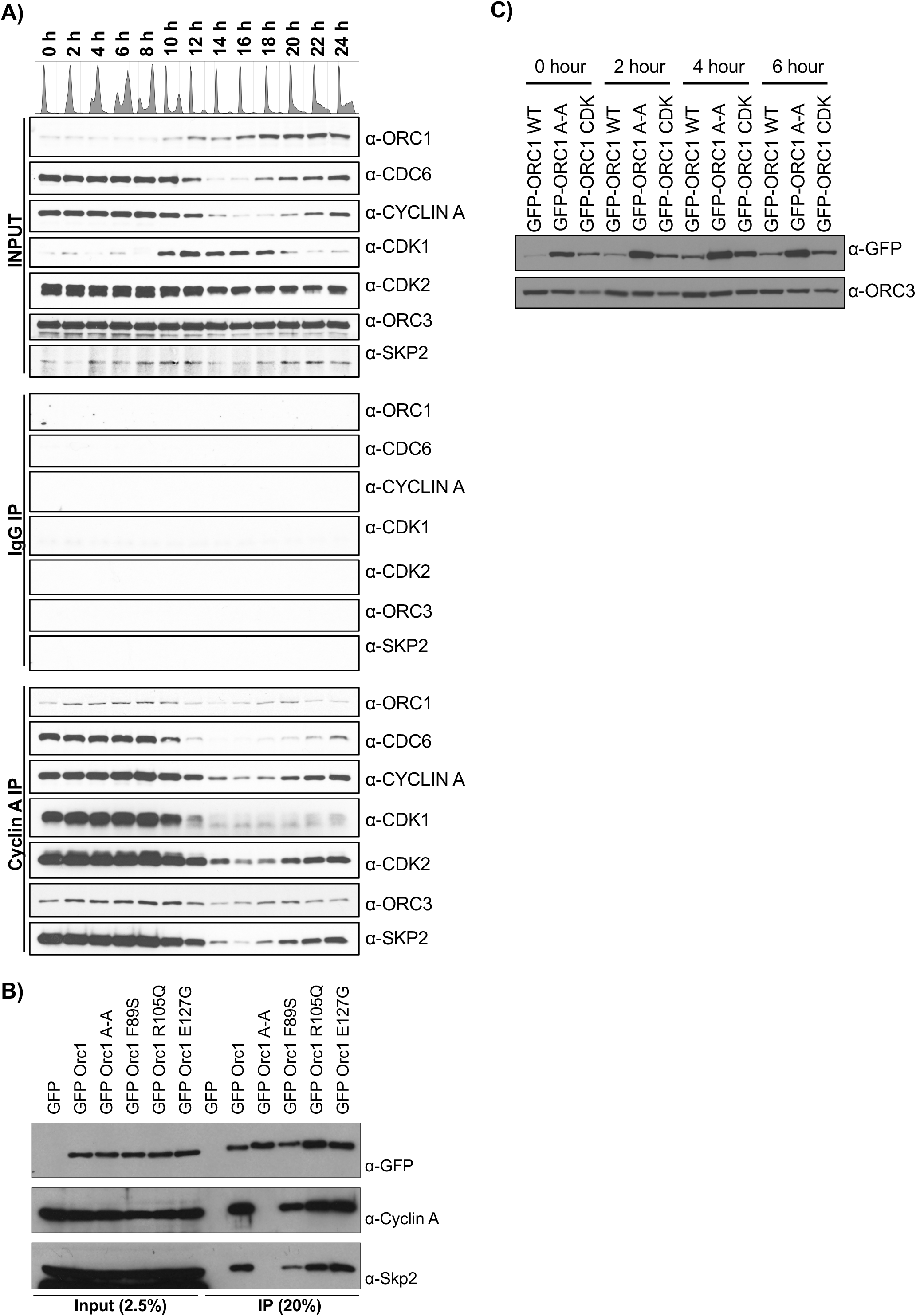
SKP2 regulates ORC1 degradation through Cyclin A binding to the ORC1 Cy motif, Related to Figure 5. (A) HeLa cells were synchronized at G1/S using double thymidine block and cells harvested at indicated time points after release from block were lysed. The whole cell extract was immunoprecipitated with anti-Cyclin A antibody and further immunoblotted with indicated antibodies. (B) HEK293 cells were transfected with GFP-ORC1 WT or the indicated Meier-Gorlin mutants. Whole cell lysates were used for immunoprecipitation with anti-GFP antibody. Immunoprecipitates were analyzed with indicated antibodies. (C) The GFP-ORC1 WT, A-A and CDK stable U2OS cell lines were synchronized at G1/S using a double thymidine block and harvested at different time points after release into fresh media. Total lysates were immunoblotted with anti-GFP and anti-ORC3 antibodies.

**Figure S6.**
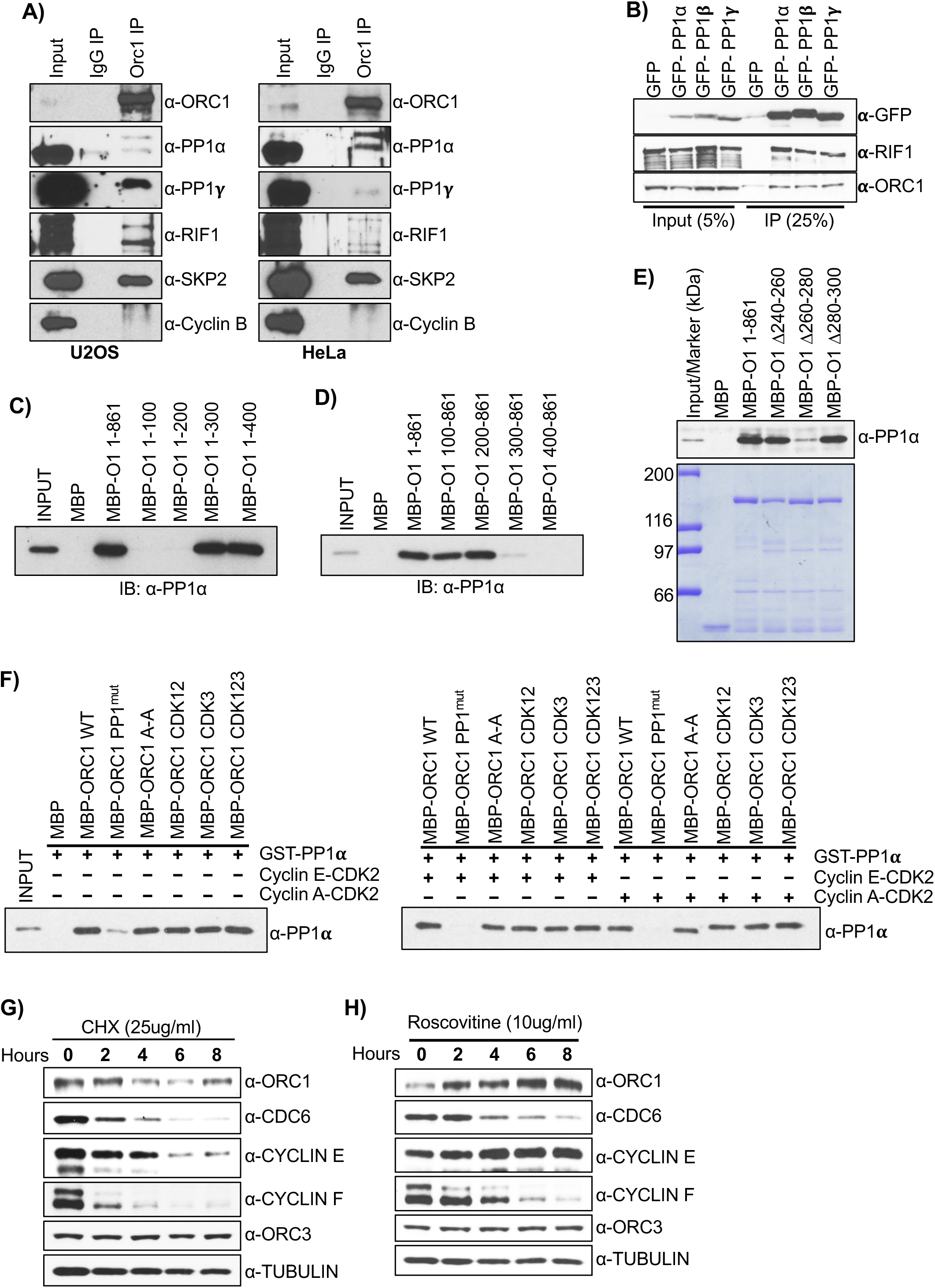
ORC1 and PP1 interaction controls the phosphorylation of ORC1 that regulates its degradation, Related to Figure 6. (A) Endogenous ORC1 was immunoprecipitated from asynchronous U2OS (left panel) and HeLa (right panel) cell lysates with anti-ORC1 mouse monoclonal antibody. The immunoprecipitates were immunoblotted with indicated antibodies. (B) Cell lysates of HEK293 cells overexpressing GFP-PP1 isoforms (α, β and γ) were used for immunoprecipitation with anti-GFP antibody and immunoblotted with indicated antibodies. (C-D) Mapping of PP1 interaction region in ORC1 protein. MBP-ORC1 WT or its C-terminal deletion mutant proteins (C) or N-terminal deletion mutant proteins (D) bound to α-MBP antibody magnetic beads were further incubated with purified recombinant GST-PP1α protein in MBP pull-down assays and immunoblotted with anti-PP1α antibody. MBP protein served as negative control. (E) The recombinant MBP-ORC1 full length as well as its various internal deletion mutant proteins as indicated, bound to α-MBP antibody magnetic beads, were incubated with purified GST-PP1α protein in MBP pull-down assays and blotted with anti-PP1α antibody (top panel). The internal regions of MBP-ORC1 protein is deleted in context of full length. Loading of MBP-fused proteins used in pull-down assay is shown in bottom panel. (F) In MBP pull-down assays, MBP-ORC1 WT or its mutant proteins as indicated, bound to α-MBP antibody magnetic beads were incubated with GST-PP1α either alone or in presence of Cyclin E-CDK2 or Cyclin A-CDK2 kinases with 1mM ATP. The western blot is probed with anti-PP1α antibody and MBP protein served as negative control. (G) Stability analysis of endogenous proteins was performed in U2OS cells by cycloheximide chase. Exponentially growing U2OS cells were treated with 25ug/ml of cycloheximide for indicated times. Total cell lysates were immunoblotted with for detection of endogenous levels of indicated proteins. (H) Asynchronous growing U2OS cells were treated with 10ug/ml of roscovitine for indicated times. Total cell lysates were immunoblotted with indicated antibodies.

**Figure S7.**
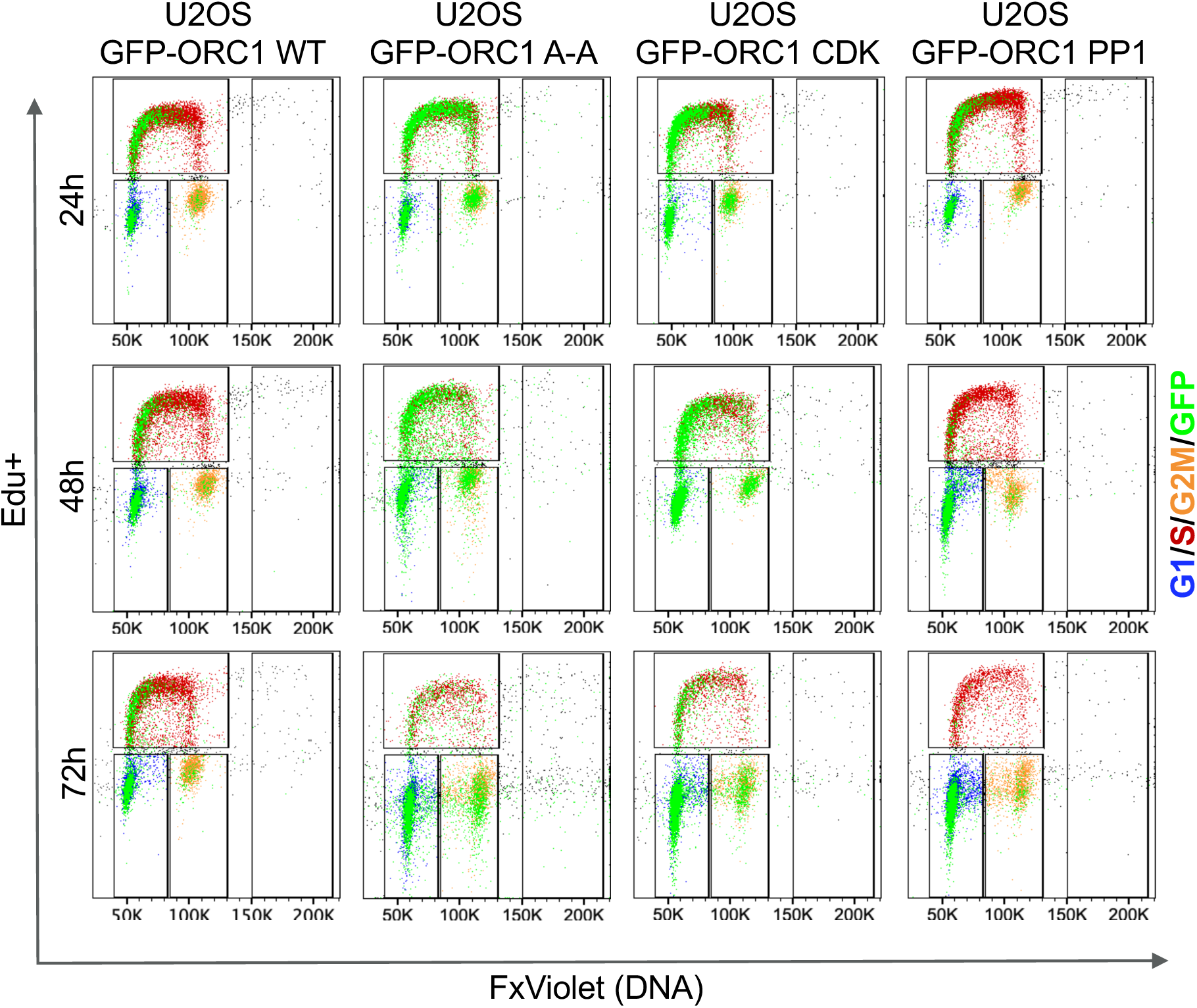
Cell cycle analysis of GFP-ORC1 wild type and GFP-ORC1 mutants. Related to Figure 7. Analytical flow cytometry profiles of GFP-ORC1 WT, GFP-ORC1 A-A, GFP-ORC1 CDK123 and GFP-ORC1 PP1mut. Cells were treated with 1 µg/ml of doxycycline and endogenous ORC1 was depleted with siRNA targeting 3’UTR of *ORC1* mRNA. Cells were pulse labeled with EdU for 2 hours prior to harvesting at 24, 48 and 72 hours. Overlay plots show expression of GFP-ORC1 (green) across cell cycle phases (G1 – blue; S – red; G2M – orange). Representative flow plots of experiment from at least 3 biological replicates.

**Figure S8.**
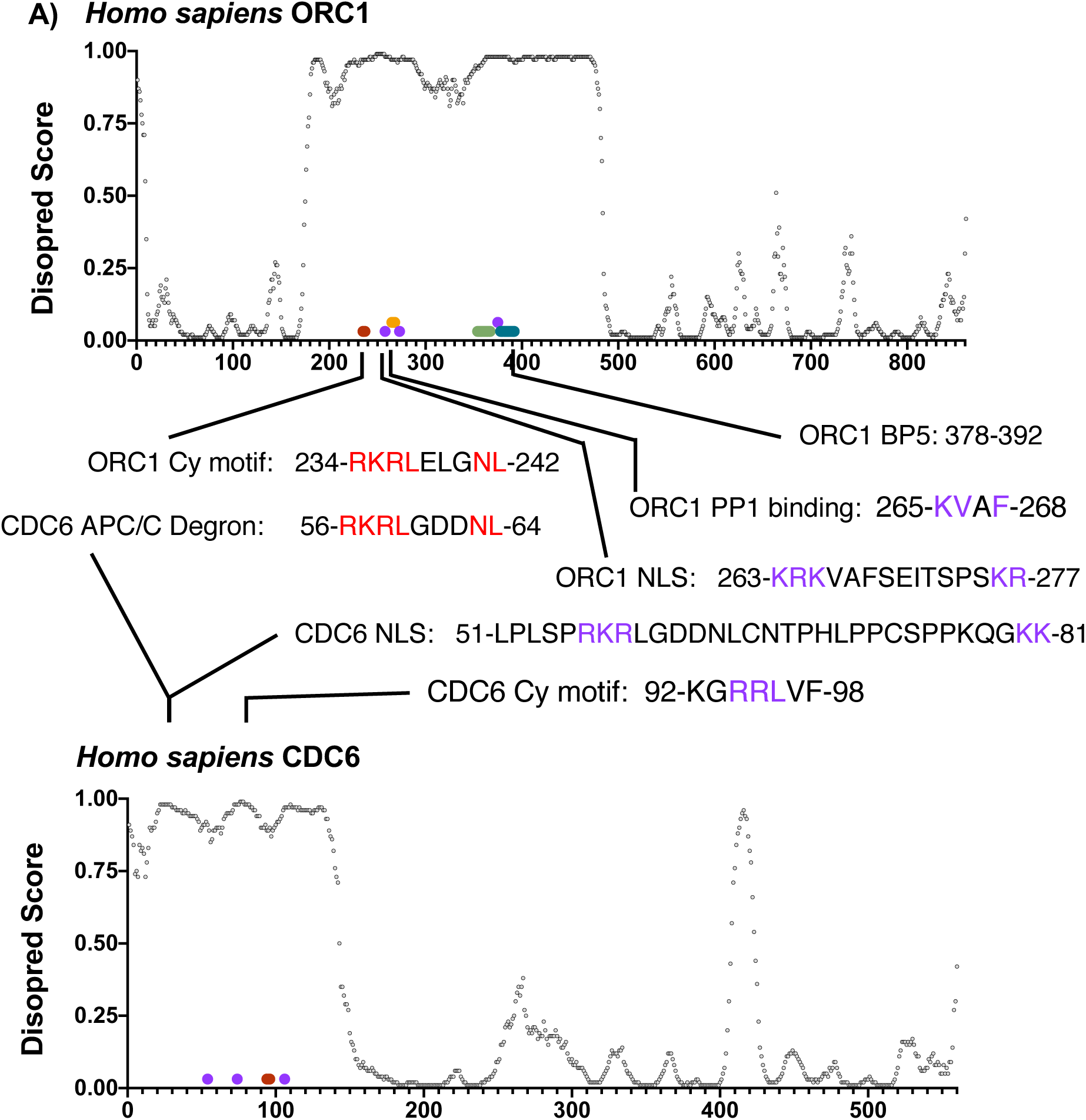
Summary of SLiMs in ORC1 and CDC6 predicted intrinsically disordered regions. The short linear peptide motifs in the ORC1 and CDC6 IDRs are shown. The similarity of the ORC1 Cy motif degron for SCF^SKP2^ and the CDC6 degron for APC/C ubiquitin ligases is shown (conserved residues in red). The CDC6 APC/C degron is from Pederson et al. (2000). The ORC1 Cy motif also functions as an ORC1 degron at the G1/S phase transition. The other motifs are shown and the consensus residues are shown in purple. The CDC6 NLS from different reports indicates that the region to vary from 54-81 aa (Delmolino et al., 2001; Takei et al., 1999).

## STAR METHODS

### CONTACT FOR REAGENT AND RESOURCE SHARING

Further information and requests for resources and reagents should be directed to and will be fulfilled by the Lead Contact, Bruce Stillman.

### EXPERIMENTAL MODEL AND SUBJECT DETAILS

#### Cell culture and cell synchronization, with siRNA treatment or transient transfection of expression plasmids

U2OS, HeLa and HEK293 cells were obtained from the Cold Spring Harbor Laboratory cell culture collection and cultured in DMEM containing high glucose (Gibco) supplemented with 10% inactivated fetal calf serum and Penicillin/Streptomycin. HeLa suspension cells were grown in suspension in Joklik Modified Eagle’s Essential Minimal Medium (US Biologicals, M3867) supplemented with 5% calf serum (Hyclone, SH30087.04). All the cell lines tested negative for mycoplasma contamination. HeLa suspension cells were synchronized by double thymidine block and release as described previously (Siddiqui and Stillman, 2007). Briefly, cells were synchronized with 2.5mM Thymidine (Sigma, T1895-25G) for 16 hours and then released into fresh complete JMEM for 12 hours. Cells were again treated with 2.5mM Thymidine for 16 hours after which, cells released into fresh media were harvested as 0 hour (at G1/S boundary) and every two hours thereafter. To synchronize U2OS cells at G2/M boundary, 100 ng/mL of nocodazole was added to fresh medium for 16 hours. After 16 hours of block the cells were washed two times with 1x Phosphate Buffered Saline (PBS) and subsequently, released into the fresh media. Mitotic U2OS cells were collected by mitotic shake-off method. U2OS cells were synchronized at G1/S using double thymidine block and release protocol. Briefly, the U2OS cells were blocked for 16 hours with incubation of 2.5mM of thymidine, washed with PBS to release for 12 hours in drug free medium and again incubated with thymidine for 16 hours. The cells were washed and released after second thymidine block for different times. For depletion of proteins in G1/S synchronized U2OS cells, the U2OS cells transiently transfected with 100nM of siRNA’s (control GFP as well as SKP2 siRNA) using Lipofectamine RNAiMax after first thymidine block. Plasmids expressing exogenous genes were transfected using 5 μg of DNA using lipofectamine 2000 transfection reagents (Thermo Fisher Scientific). The U2OS cells were incubated 25 ug/ml of cycloheximide and 10 ug/ml of roscovitine drugs as indicated in the results. The sequences of the siRNAs used are listed in key resource table.

#### Characterization of stable ORC1 mutants in U2OS cells

Wild type GFP-ORC1 as well as GFP-ORC1 A-A, GFP-ORC1 CDK and GFP-ORC1 PP1 mutants were cloned into the plasmid pcDNA ™5/FRT/TO (Life Technologies) under a tetracycline-regulated CMV-based promoter. The plasmids were integrated into a FRT-U2OS-TRex osteosarcoma cells (Malecki et al., 2006) via FRT-mediated recombination and the integrated transgenes were selected with hygromycin. The expression of ORC1 was assessed following treatment of cells with 1 ug/ml of Doxycycline.

### METHOD DETAILS

#### Plasmid Construction and Mutagenesis

Recombinant human MBP tagged ORC1 protein or its mutants, ORC2, ORC3, ORC4, ORC5, ORC6 and CDC6 were expressed and purified from bacteria by cloning into pMALp-c2E vector. MBP-ORC1 180-240 WT and A-A were also cloned in pMALp-c2E vector with an addition of prescission protease site at N-terminus and 6XHis tag at C-terminus. The human ORC1 WT or its mutants were also cloned in pEGFP-C1 vector for transient over-expression in mammalian cells. Human CDC6 or its mutants and PP1α were also cloned in pGEX-6P1 vector to produce GST fusion proteins in bacterial cells. PP1α plasmids were also generated with a N-terminal T7 tag in the pLPC vector (McCurrach et al., 1997) (gift from Scott Lowe, Memorial Sloan Kettering Cancer Center) for mammalian expression.

The mutant plasmids were generated following site directed mutagenesis protocol using Phusion high fidelity DNA polymerase (NEB). To generate internal deletions of MBP-ORC1 and GST-CDC6 plasmids, we employed phosphorothioate-modified PCR primers and T7 gene 6 exonuclease following a protocol previously described (Stoynova et al., 2004). All of the plasmid constructs were verified by sequencing. Oligonucleotide sequences used to generate the plasmids are available upon request.

#### Expression and purification of recombinant proteins

The MBP- and GST-fusion recombinant proteins were expressed and purified using amylose and Glutathione beads, respectively following a previously described protocol (Hossain and Stillman, 2012). Briefly, the MBP and GST fusion were transformed into E. coli BL21 cells with their respective plasmids, transformed bacterial cells were grown in LB media at 37°C till the O.D. of the cells reaches 0.7-0.9 and were induced for 12 hours with 0.3mM of IPTG at 16°C. The induced cells were pelleted, washed, and further lysed with sonication in a lysis buffer A containing 25mM Tris-HCl at pH 7.5, 150mM NaCl, 0.02% NP-40, 5mM benzamidine-HCl, 1mM phenylmethylsulfonylfluoride, Protease cocktail inhibitor tablets [Roche], 10% glycerol) plus 100mg/ml lysozyme. The lysed bacterial cells were centrifuged and the clarified supernatant is incubated with respective pre-washed Amylose or Glutathione agarose beads for 3 hours at 4°C. The bead bound proteins were washed with ten column volumes of buffer A plus 0.05% NP-40 + 500mM NaCl and further with ten column volumes of buffer A alone. Fusion protein was eluted in a stepwise manner with buffer A containing 20mM Maltose for MBP fusion proteins or 20 mM reduced glutathione, pH7.5 for GST fusion proteins. Fractions containing purified proteins were pooled, concentrated and dialyzed, and protein concentration was estimated using a standard Bradford protein assay. His-tagged ORC1 180-240 WT and A-A mutant were made after cleaving MBP protein from the N-terminus using prescission protease and further purified using Ni-NTA resin (Qiagen).

Cyclin A-CDK2 and Cyclin E-CDK2 kinases were expressed and purified from baculoviral infected insect cells as described previously (Hossain and Stillman, 2012).

#### MBP and GST pull-down assay

For MBP and GST pull-down assay, anti-MBP or anti-GST antibodies were initially bound to gamma bind protein G magnetic beads, washed and further incubated with respective purified recombinant MBP- and GST-fusion proteins in binding buffer with following composition; 25mM Tris-Cl at pH 7.5, 150mM KCl, 0.15% Nonidet P-40, 0.1mM EDTA, 5 mM magnesium acetate, 1mM DTT. For MBP-ORC1 pull down, MBP-ORC1 protein bound beads were either incubated with GST-CDC6, GST-PP1α, Cyclin E-CDK2, Cyclin A-CDK2 or ORC1 180-240-His depending on context of experiments for 4-5 hours at 4°C. For GST-CDC6 pull down, the GST-CDC6 protein bound magnetic beads were either incubated MBP-ORC1 alone or in combination with Cyclin E-CDK2 or/and Cyclin A-CDK2 in the presence or absence of 1mM ATP. The beads were washed with binding buffer and was further immunoblotted with respective antibodies as indicated in the results.

#### Immunoprecipitation

For expression of proteins, HEK293 cells were transiently transfected with the indicated plasmids with lipofectamine 2000 transfection reagent (ThermoFisher Scientific). For immunoprecipitation, the cells were harvested and washed in PBS and lysed in a buffer containing 20mM Tris-HCl pH7.5, 300mM NaCl, 0.3% NP-40, 5mM MgCl_2_, 0.1 mM EDTA, 10% Glycerol, 1mM DTT, 1mM CaCl_2_, 20uM MG132 and protease as well as phosphatase inhibitor tablets (Roche). Benzonase (Sigma-Aldrich, St. Louis, MO) was added to the buffer and the suspension incubated for 30 min on ice with intermittent mixing. The concentration of NaCl and NP-40 was reduced to 150mM and 0.15%, respectively with dilution buffer after 30 minutes incubation on ice. The extract was centrifuged at 14, 000 rpm for 15 min at 4°C. The GFP-ORC1 and GFP-PP1 proteins were precipitated with anti-GFP specific antibodies as indicated in figure legends. The whole cell extract was first incubated with antibodies for 4 hours and subsequently, 2 hours with pre-washed gamma bind G sepharose beads with end-to end shaking at 4°C. The beads were washed 3 times with washing buffer containing 20mM Tris-HCl pH7.5, 150mM NaCl, 0.15% NP-40, 5mM MgCl_2_, 0.1mM EDTA, 10% Glycerol, 1mM DTT and protease as well as phosphatase inhibitor tablets from Roche. Finally, the washed beads were suspended in Laemmli sample buffer and 8% SDS-PAGE gels were run and immunoblotted.

For co-immunoprecipitation of ORC1, CDC6 and Cyclin A interacting proteins, extracts were prepared by resuspending the synchronized and asynchronous cells in Lysis Buffer A (20mM Tris-HCl pH 7.5, 250mM NaCl, 5mM MgCl2, 0.1mM EDTA, 0.3% NP40, 1mM DTT, 1mM CaCl2, 5% Glycerol supplemented with Benzonase, Roscovitine, MG132, protease inhibitor cocktail and phosphatase inhibitors) for 30 minutes on rotation at 4°C. Following lysis, the extract was diluted with equal volume of Dilution Buffer B (Buffer A without NaCl and NP-40) to adjust the final salt and detergent concentration to 125mM NaCl and 0.15% NP-40 respectively, and centrifuged to obtain clarified extract. Half of each extract was used to perform immunoprecipitations using either CDC6 (CS1881) or ORC1 antibody and corresponding isotype control bound to protein G dynabeads. The beads were washed and were then suspended in 2X Laemmli sample loading buffer, and analyzed by western blot.

For immunoprecipitations, the following antibodies were used: mouse monoclonal anti-GFP antibody (A11120; Invitrogen), rabbit polyclonal anti-CDC6 antibody (CS1881, CSHL facility), rabbit polyclonal anti-Cyclin A antibody (A305-254A; Bethyl), mouse monoclonal ORC1 78-1-172 (Kara et al., 2015) and mouse monoclonal anti-T7 antibody (CSHL Facility). For immunoblotting, the antibodies are indicated in the key resource table under antibodies section.

#### EdU incorporation and Cell cycle analyses of mutant ORC1

U2OS cells were cultured in DMEM containing high glucose (Gibco) supplemented with Penicillin/Streptomycin and 10% fetal bovine serum, or in case of tetracycline-inducible U2OS cells stably expressing GFP-ORC1 WT, GFP-ORC1 A-A, GFP-ORC1 CDK123 and GFP-ORC1 PP1^mut^ in 10% Tet-free fetal bovine serum (Takara, 631101) and 150 ug/ml Hygromycin. On day 0 all cells were induced with 1 µg/ml doxycycline (Sigma, D9891-10G) and thereafter maintained in it for the remainder of the experiment. Cell lines were treated on day 0 with (a) 200nM of either control or 3’UTR ORC1 siRNA to deplete endogenous ORC1 to be collected 24 hours later, and (b) 100nM of either control or 3’UTR ORC1 siRNA on day 0 and again on day 1 (24 h), for samples to be collected on day 2 (48 h) and day 3 (72 h). All samples were treated with 10µM EdU for 2 hours prior to harvesting. Samples were either counted for growth curves, or prepared for cell cycle analyses using the Click-iT Plus EdU Alexa Fluor 647 flow cytometry assay kit (Invitrogen, C10635) as per the manufacturer’s protocol. Three-color flow cytometric analysis was performed using a BD LSRFortessa Dual SORP (BD Biosciences, San Jose, CA). The GFP emission was collected off the 488nm laser using a 505LP dicroic mirror and a 530/30 filter, Alexafluor 647 emission was collected off the 638nm laser using a 670/30 filter and the FxViolet emission was collected off the 407nm laser using the 450/50 filter.

#### Live cell microscopy

Tetracycline-inducible GFP-Orc1 WT and GFP-ORC1 A-A stable U2OS cell lines were used for live-cell imaging after addition of 1 ug/ml doxycycline. The cells were transferred to a 6-well plates mounted onto the stage of spinning disk confocal microscope (Perkin Elmer) and kept at 37°C in DMEM medium with 10% FBS. Time-lapse images acquired with a 40X objective lens were captured with a CCD camera or the Perkin Elmer spinning disk acquisition setup.

### QUANTIFICATION AND STATISTICAL ANALYSIS

Data points in figures represent mean values, with error bars in all figures represent mean ± Standard Deviation. Number of replicates are indicated in the figure legends.

